# Discovery of CDK4-selective molecular glue degraders by high-throughput proteomics

**DOI:** 10.64898/2026.06.19.733443

**Authors:** Patrick R. A. Zanon, Bachuki Shashikadze, Denise Winkler, Ines Scheller, Denis Bartoschek, Anastasia Bednarz, Tobias Graef, Sophie Machata, Uli Ohmayer, Bjoern Schwalb, Martin Steger, Henrik Daub

## Abstract

Molecular glue degraders (MGDs) are proximity-inducing molecules that promote the destruction of disease-causing proteins by stabilizing novel interfaces between E3 ubiquitin ligases and target proteins. The rational design of MGDs remains exceptionally challenging, historically relying on serendipitous discoveries. Here, we deployed a high-throughput, mass spectrometry (MS)-based screen evaluating thousands of cereblon (CRBN)-directed compounds to expedite the identification of novel neosubstrates. This workflow led to the discovery of **NE26394**, a first-in-class MGD that selectively eliminates cyclin-dependent kinase 4 (CDK4), a critical oncogenic driver of cell cycle progression. Mechanistically, **NE26394**-induced CDK4 recognition by CRBN depends on the co-recruitment of endogenous INK4 family proteins. In CDK4-dependent cancer models, **NE26394** effectively mimics the anti-proliferative RB-E2F pathway perturbations induced by clinical CDK4 inhibitors, rendering it an attractive candidate for further preclinical development.

## Introduction

The eukaryotic cell cycle encompasses a series of precisely orchestrated cellular processes that are tightly controlled by cyclin-dependent kinases (CDKs) and their complex partners, the cyclins. Transition from G1 into S phase of the cell cycle is primed by proliferative signals that induce expression of D-type cyclins and the activation of CDK4 and CDK6.^1,2^ CDK4/6-mediated phosphorylation of the tumor suppressor retinoblastoma protein (RB) and its related family members leads to the release of E2F transcription factors and, consequently, de-repression of a large gene set required for cell cycle progression.^3^ This also triggers cyclin E-dependent activation of CDK2, which induces hyperphosphorylation and complete inhibition of RB to promote full E2F-dependent transcription and cellular entry into S phase.^4^ Deregulation of the RB pathway frequently occurs in human cancers due to mutational RB inactivation or aberrant CDK4/6 activity. The latter can result from amplification, overexpression, or mutation of either CDK4, CDK6 or their companion D-type cyclins, or, alternatively, from loss-of-function of INK4 (inhibitor of CDK4) proteins.^5^ INK4 family proteins directly bind to CDK4 and CDK6, inhibiting their catalytic activity and preventing them from pairing with D-type cyclins. In humans, they are encoded by four related genes, CDKN2A, CDKN2B, CDKN2C and CDKN2D, encoding the INK4 family proteins INK4A (p16), INK4B (p15), INK4C (p18) and INK4D (p19), respectively. Despite their shared function in inhibiting CDK4/6 activity, mainly CDKN2A is considered a major tumor suppressor gene, as it exhibits the highest mutation frequency.^6,7^

The relevance of the RB pathway has spurred the development of small-molecule kinase inhibitors targeting CDK4 and CDK6 to restrict tumor cell proliferation. These efforts culminated in the demonstration of clinical efficacy in combination with endocrine therapy, followed by the approval of the CDK4/6-selective drugs palbociclib, ribociclib and abemaciclib for first-line therapy in hormone receptor-positive, HER2-negative breast cancers.^8^ Notably, suppression of CDK4 activity is the key determinant of sensitivity in cancer cell lines harboring no or little CDK6 protein. In contrast, most CDK6-expressing cell lines exhibit intrinsic CDK4/6 inhibitor (CDK4/6i) resistance despite efficient *in vitro* inhibition of CDK6-cyclin D3 kinase activity.^4^ Investigation of the underlying mechanisms revealed attenuated CDK4/6i binding to CDK6 in resistant tumor cells, as the kinase adopted a thermostable conformation interacting weakly with the HSP90-CDC37 chaperone when compared with drug-sensitive cellular backgrounds.^9^ Depending on cellular context, this resilience to CDK4/6i may be induced upon binding of INK4 proteins, as partial preservation of CDK6 activity has been reported upon formation of INK4C-cyclin D-CDK6 complexes.^10^ In contrast, CDK6 activity is efficiently suppressed in hematological cancer cell lines and in immune cells like CD34^+^ hematopoietic stem and progenitor cells or mature neutrophils. This represents a major liability, causing neutropenia in CDK4/6i-treated patients and resulting in dose-limiting toxicity.^11^ Collectively, these data highlight CDK4 as the key efficacy target for CDK4/6i therapy, showing clinical benefit in tumors that predominantly express CDK4 and depend on its activity, while dose-limiting hematological adverse events are primarily due to CDK6 inhibition.^11^ A recent landmark study by Anders and colleagues reported impressive preclinical anti-tumor efficacy and safety data for the next-generation CDK4-selective, largely CDK6-sparing inhibitor atirmociclib,^12,13^ which is currently being evaluated in several clinical trials in HR+ breast cancer.^14^

More recently, heterobifunctional degraders (or proteolysis-targeting chimeras (PROTACs)) hijacking CRL4^CRBN^ E3 ubiquitin ligase have been reported that induce degradation of either CDK4, CDK6, or both protein kinases simultaneously.^10,15,16^ Such targeted protein degradation (TPD) approaches embrace the groundbreaking paradigm of eliminating rather than inhibiting disease-causing proteins to achieve complete, prolonged, and highly selective target inactivation.^17,18^ In addition to PROTACs, molecular glue degraders (MGDs) have emerged as major TPD modality that engage E3 ligase substrate receptors like CRBN. CRBN-based MGDs are smaller in size than PROTACs and act by creating complementary neomorphic surfaces, triggering CRBN recruitment of non-native E3 ligase substrates (neosubstrates) followed by CRBN-mediated ubiquitination and subsequent proteasomal degradation.^17,19^ CRBN-targeted MGDs of the transcription factors IKZF1/3, such as the immunomodulatory imide drug (IMiD) lenalidomide, have been used in multiple myeloma treatment for two decades^20,21^ and various other neosubstrates were discovered more recently.^22–25^ Intriguingly, the number of neosubstrates is now growing exponentially owing to the advent and massive impact of high-throughput proteomics screening platforms.^26–30^ We previously reported a 96-well plate based cellular screening format comprising highly parallelized sample preparation, single-shot data-independent acquisition (DIA) mass spectrometry (MS) and automated data analysis. This workflow enabled proteome-wide interrogation of degrader selectivity and efficacy through quantitative profiling of up to 11,000 proteins.^26,30^ Follow-up experiments confirmed CRBN- and cullin RING (CRL) E3 ligase-dependent degradation for more than 200 novel CRBN neosubstrates that were mechanistically validated in molecular glue-induced proximity and cellular ubiquitination assays, emphasizing the broad target scope of the MGD drug discovery modality. Remarkably, our earlier screens recorded many neosubstrate candidates for only one or two compounds out of about 1,000 tested CRBN-binders.^30^ This finding likely reflects the steep structure-activity relationships (SARs) governing ternary complex formation and productive protein degradation. Interestingly, in addition to many zinc finger proteins harboring a β-hairpin glycine loop structural degron motif for CRBN recognition, protein kinases appear to constitute a prominent subset of CRBN MGD targets, as exemplified by recent reports on *e.g.* WEE1,^31^ NEK7,^32^ CDK2,^33^ TBK1,^29^ IRAK1 or AKT3.^30^

Here, we report CDK4 as a novel CRBN MGD target, discovered through an unbiased, large-scale proteomics screening campaign spanning several thousand compounds. Chemical optimization resulted in a highly efficacious, potent, and selective CDK4 degrader inhibiting the RB-E2F pathway in CDK4-dependent cancer cells.

## Results

### Discovery of a CDK4-degrading molecular glue compound by integrated proteomic screening and neosubstrate validation

High-throughput proteomics of CRBN-directed compound libraries has emerged as a powerful strategy for the scalable discovery of novel MGDs and their cellular targets (Fig. 1a).^26,27,29,30^ When screening CRBN-based MGDs derived from a phenyl-glutarimide CRBN warhead,^34^ we previously encountered neosubstrate profiles that were strikingly different from those of MGDs harboring classical lenalidomide or thalidomide core scaffolds.^26^ This key finding provided a strong rationale to further expand proteomics screening to CRBN-directed compound collections built on chemically diverse scaffolds. Our aim was to systematically map the MGD-accessible target space for disease-relevant proteins currently deemed undruggable or inadequately targeted by classical small molecule pharmacology. To achieve this, we leveraged unbiased proteomic profiling across a library of several thousand molecules in CRBN-overexpressing (CRBN^oe^) cells to maximize neosubstrate identification^30,35^. This screening campaign led to the discovery of **NE25613**, a small molecule that depleted CDK4 levels by more than 90% within 6 h of treatment (Fig. 1b). Co-treatment with either the NEDD8-activating enzyme inhibitor MLN4924 or the proteasome inhibitor bortezomib completely suppressed **NE25613**-induced CDK4 downregulation, establishing CRL- and proteasome-dependent mechanisms (Fig. 1b). The second-most pronounced, MLN4924- and bortezomib-dependent downregulation occurred for the nuclear pre-mRNA domain-containing protein 1B (RPRD1B), a transcriptional regulator known to interact with the phosphorylated C-terminal domain of RNA polymerase II. It modulates alternative polyadenylation of cell growth transcripts, with potential relevance to cancer cell biology.^36,37^ Comparison of **NE25613**-triggered protein changes in the absence and presence of MLN4924 additionally revealed CRL-dependent regulation of the RPRD1B-interacting proteins RPRD1A and RPRD2 (Fig. 1c). In contrast to these MLN4924-sensitive targets, neddylation inhibitor treatment failed to block the downregulation of many other proteins, including PCLAF, the mitotic kinases AURKA and PLK1, and, remarkably, even established CDK4 complex partners such as cyclin D1 (CCND1) or p21/CIP1 (CDKN1A). We have previously observed similar off-target profiles for certain CRBN MGDs exhibiting high compound activity in global proteomics experiments.^26,30^

**Fig. 1:**
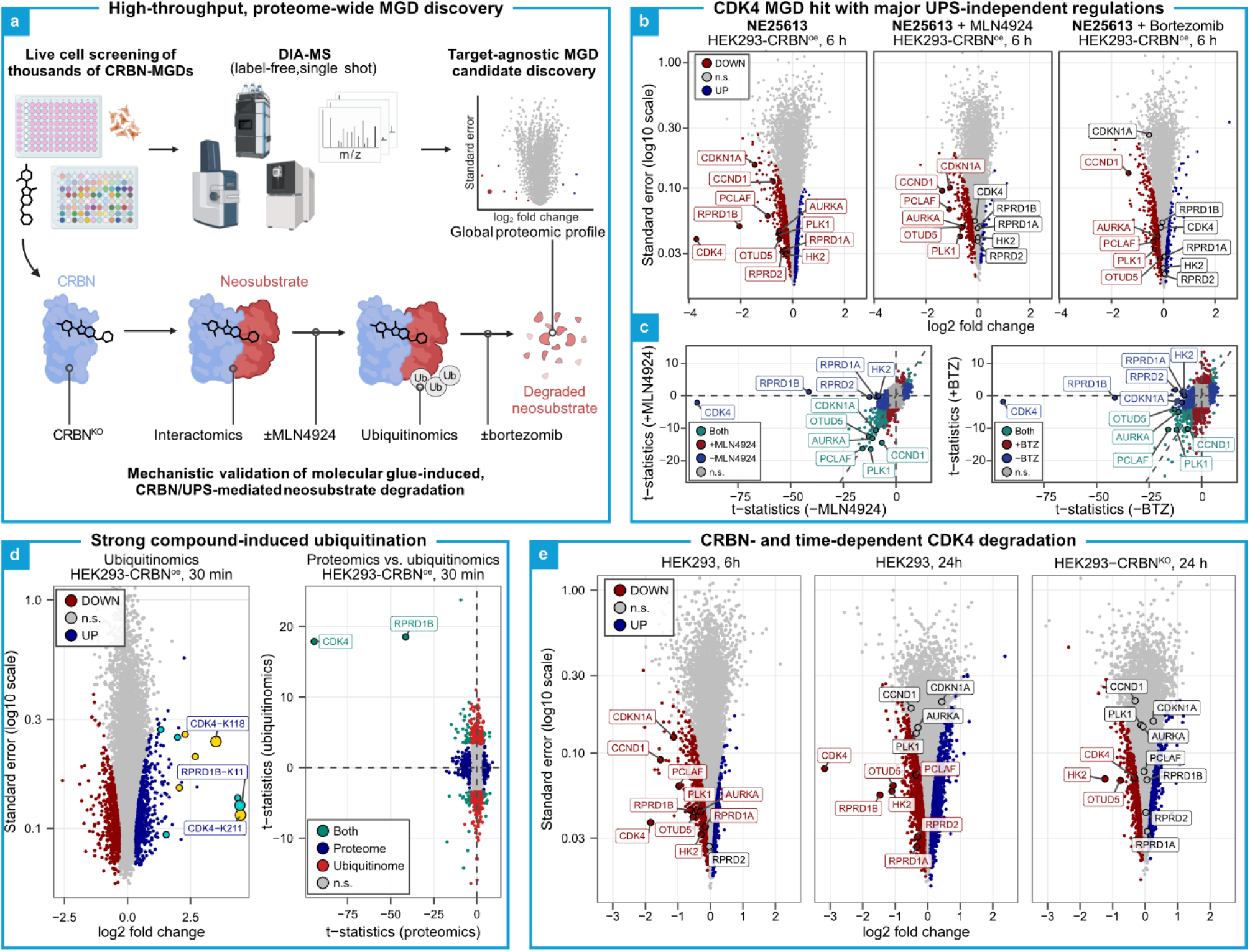
Identification and validation of the CDK4 as molecular glue degrader target of NE25613. **a**, Proteome-wide screening of CRBN molecular glue libraries using high-throughput proteomics and validation of degrader mode-of-action, which encompasses counter-screening in a CRBN-knockout cell line (CRBN^KO^), demonstrating of induced proximity of the neosubstrate to CRBN (interactomics), validating dependency on functional cullin-RING E3 ligases (CRL) (±MLN4924), showing ubiquitination of the neosubstrate (ubiquitinomics), and verifying proteasomal dependency (±bortezomib). **b**, Volcano plot visualisations (log_2_ fold-change (x-axis) versus log_10_ standard error (y-axis) of the proteome regulations in HEK293-CRBN^oe^ cells treated for 6 h with **NE25613** alone (left), or co-treated with MLN4924 (centre), or bortezomib (right). Selected proteins are labelled. **c**, Comparison of the t-statistics for **NE25613**-treated HEK293-CRBN^oe^ cells co-treated with DMSO (x-axis), MLN4924 (y-axis, left) or bortezomib (y-axis, right). Statistically significant up- and downregulations (FDR=1%) are colored in blue and red, respectively. Proteins that were significantly downregulated in both shown conditions are highlighted in turquoise. n.s.= not significant. **d**, Site-specific identification of differentially regulated ubiquitination events through K-GG remnant profiling (left). On the right, a comparison of the t-statistics (**NE25613** vs. DMSO) of the proteome (x-axis) and the ubiquitinome (y-axis) is shown. **e**, Volcano plot visualisation of proteomic profiles of **NE25613**-treated parental HEK293 cells (6 h, left), 24 h (centre), and in HEK293-CRBN^KO^ cells after 24 h (right). In volcano plots, log_2_ fold changes relative to vehicle (x-axis) and standard errors (y-axis, log_10_ scale) are shown. For MLN4924 and bortezomib co-treatments, fold-changes relative to vehicle + the respective inhibitor are shown. Up- and downregulated proteins (mod. adj. p-value < 0.01) or ubiquitination sites (adj. p-value < 0.05) are coloured in blue and red, respectively. Non-significant (n.s.) regulations are coloured grey. In comparison plots, the t-statistics for regulations in two different experiments are plotted against each other. For comparison of global proteomics and ubiquitinomics profiles, ubiquitination sites mapping to the same protein were averaged, and only significantly upregulated sites were included in the analysis. Statistically significant up- and downregulations in the x- and y-axis experiments are coloured in blue and red, respectively, overlapping significant features are coloured turquoise, and non-significant regulations are coloured grey.

To mechanistically characterize the CRL-dependent downregulations, we next performed global ubiquitinomics in HEK293-CRBN^oe^ cells following an acute 30-min treatment with **NE25613** (Fig. 1d). After cell lysis, di-Gly (or K-GG) remnant peptides harbouring cellular lysine ubiquitination sites were immunopurified and subsequently analysed *via* single-shot DIA-MS.^38^ We detected multiple strongly induced ubiquitination sites mapping to both CDK4 and RPRD1B, indicating CRBN-mediated cellular ubiquitination upon MGD-induced ternary complex formation, thereby confirming these two proteins as novel CRBN neosubstrates. (Fig. 1b-d). In addition, we recorded rapid induction of ubiquitination and MLN4924-sensitive protein downregulation for RPRD1A and RPRD2. These RPRD1B-associated factors likely represent bystander neosubstrates, such as the CSNK1A1-interacting FAM83 proteins or the CDK-activating kinase (CAK) complex members CDK7 and CCNH that are recruited to CRBN by MNAT1.^26,32^

To corroborate CDK4 degradation at physiological CRBN expression levels, we next evaluated **NE25613** by global proteomics in parental HEK293 cells (Fig. 1e). Compared to the more than 12-fold downregulation of CDK4 in the CRBN overexpression background, we observed a less pronounced, approximately 4-fold decrease in parental cells, consistent with a CRBN-mediated mechanism. We further demonstrated time-dependent CDK4 depletion in parental cells, achieving an about 90% reduction within 24 h of **NE25613** incubation, which was almost completely abrogated in CRBN knockout (CRBN^KO^) cells (Fig. 1e). While the depletion of RPRD1B and its binding partners (RPRD1A, RPRD2) was strictly CRBN-dependent, a vast majority of less pronounced regulations persisted in the CRBN^KO^ background. Because these changes were comparable to those observed in MLN4924-treated cells, we conclude that their downregulation reflects CRBN-independent off-target activity. Interestingly, a comparison of 6 h and 24 h treatment data revealed that a subset of CRBN-independent protein regulations was transient (*e.g.*, AURKA, PLK1, CCND1 or CDKN1A) whereas others intensified over time (*e.g.*, HK2 or OTUD5). These divergent dynamics prompted us to investigate whether rational chemical optimization of **NE25613** could decouple these off-target effects from on-target activity, and whether selectivity could be enhanced for CDK4 over RPRD1B. Encouragingly, our broader proteomics screen had identified an analogue that degraded RPRD1B, yet completely lacked CDK4 degradation activity and displayed negligible CRBN-independent off-target effects (Supplementary Fig. 1). This provided a strong structural precedent that these neosubstrates could be differentiated and that chemical optimization could shift selectivity toward CDK4.

### CDK4 MGD hit optimization towards higher efficacy, potency and selectivity

While our primary screening hit **NE25613** displayed excellent CDK4 degradation efficacy, its concomitant on- and off-target liabilities would likely confound phenotypic evaluations in disease-relevant models. The rational optimization of molecular glues remains inherently challenging, as minor structural modifications can trigger substantial shifts in degradation efficacy and neosubstrate selectivity profiles.^24,37,38^ Consequently, we synthesized a focussed library of 39 **NE25613** derivatives and leveraged global proteomics to perform proteome-wide structure activity relationship (SAR) analysis, a strategy uniquely suited to map the impact of structural compound modifications across the global proteome.^26,30^

In line with a recently proposed simplified model for rationalized MGD design,^17^ we divided the compound structure in three parts: (1) the glutarimide core scaffold engaging the thalidomide-binding pocket (TBP) of CRBN; (2) the proximal region up to ∼9.5 Å from the TBP core driving neosubstrate selectivity through interactions with additional CRBN residues and the degron motif; and (3) the distal shell up to ∼15 Å from the TBP core, able to further enhance the affinity for one or both of the engaged proteins (Fig. 2a).^17,23^

**Fig. 2:**
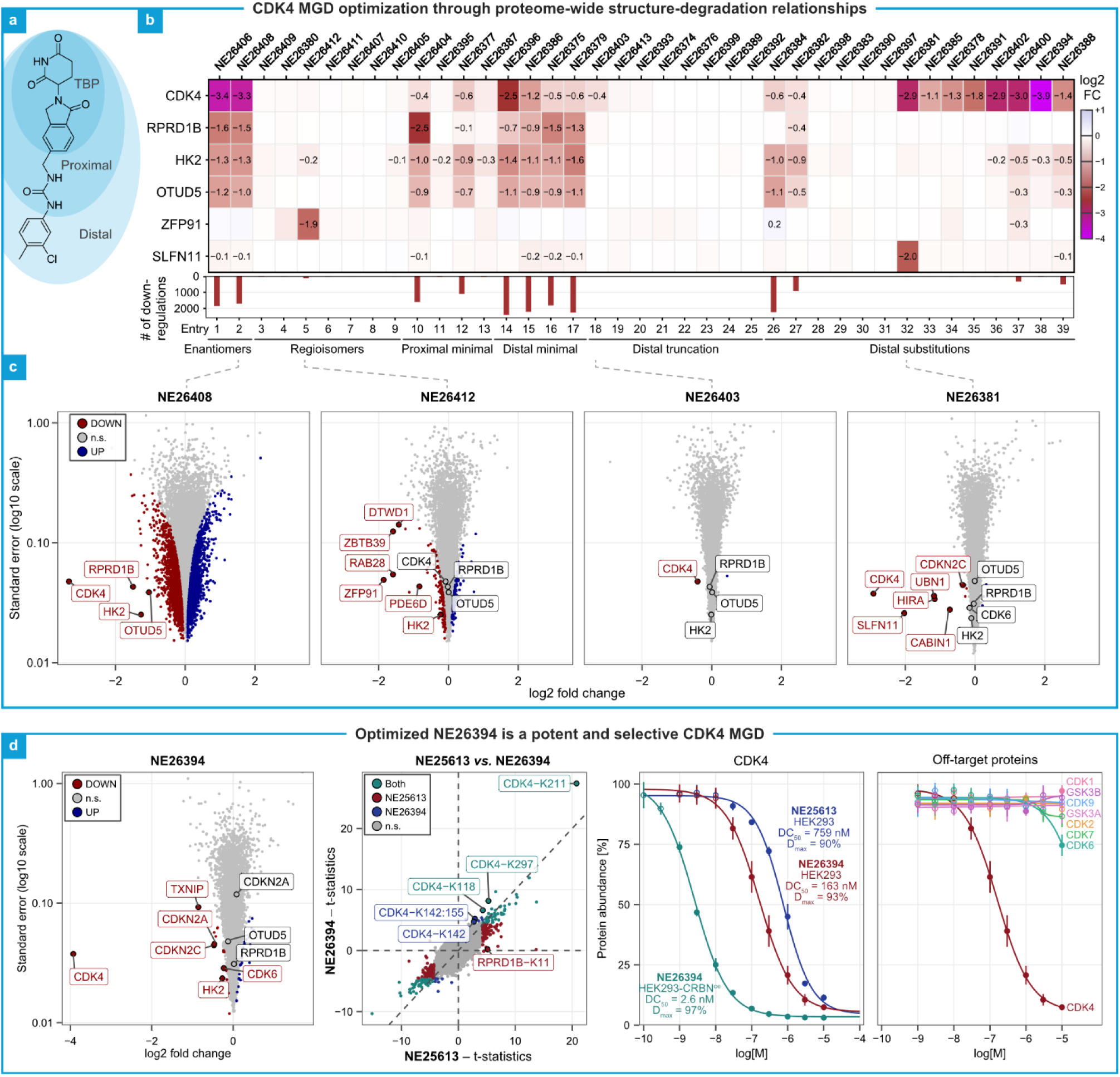
Systematic structure-activity relationships of CDK4 MGDs in HEK293 cells. **a**, Illustrative example overlaying the 2D structure of the GSPT1 MGD CC-885 with three regions of interaction within the ternary complex, as proposed by Thomä and colleagues.^17^ **b-d**, Regulation of CDK4 abundance and other prominently depleted proteins following treatment of HEK293 cells at 10 µM for 24 h. **b**, While the heatmap (top) provides individual log_2_ fold changes, the bar graph (bottom) displays the number of statistically significant downregulations emblematic of broader proteomic perturbation. **c,d**, Global proteomic regulations of selected analogues highlighting disparate target profiles and the identification of the optimized CDK4 MGD **NE26394**. Log_2_ fold-changes relative to vehicle (x-axis) and standard errors (y-axis, log_10_ scale) are shown. Up- and downregulated proteins (mod. adj. p-value < 0.01) are coloured in blue and red, respectively. Non-significant (n.s.) regulations are coloured grey. **e-g**, Further proteomic characterization of the most potent CDK4 MGD analogue. Treatment of HEK293 cells (10 µM, 30 min) with analogue **NE26394** induces stronger and ubiquitination of CDK4 than the initial hit compound **NE25613** (**e**). Potent and efficient CDK4 degradation in HEK293 and HEK293-CBRN^oe^ (**f**) while sparing other CDK family members and GSK3α/β (**g**). In volcano plots, log_2_ fold changes relative to vehicle (x-axis) and standard errors (y-axis, log_10_ scale) are shown. Up- and downregulated proteins (mod. adj. p-value < 0.01) or ubiquitination sites (mod. adj. p-value < 0.05) are coloured in blue and red, respectively. Non-significant (n.s.) regulations are coloured grey. In the comparison plot, the t-statistics for regulations in two different ubiquitinomics experiments are plotted against each other. Statistically significant up- and downregulations in the x- and y-axis experiments are coloured in blue and red, respectively, overlapping significant features are coloured turquoise, and non-significant regulations are coloured grey.

While the preference of glutarimide stereochemistry varies for CRBN MGDs, both enantiomers may engage the TBP and effect ternary complex formation.^39^ Like most glutarimids, **NE25613** was synthesized as a racemic mixture, as the stereocenter frequently is unstable and readily epimerizes in solution.^34,40,41^ In line with this, no differences in proteomic regulations were observed between the stereoisomers **NE26406** and **NE26408** (Fig. 2b,c, Supplementary Fig. S2). We therefore proceeded testing racemic mixtures. Beyond binding CRBN, a critical function of the core scaffold is to set up the vector for the rest of the molecule.^17^ As the ternary complex may accommodate different geometries, we wanted to ensure that the arrangement in **NE25613** is optimal for potency and selectivity. To this end, we synthesized seven regioisomers, which degraded neither CDK4 nor RPRD1B. Moreover, in contrast to **NE25613**, none of them displayed a CRBN-independent cellular response. We monitored this activity by quantifying the prominent off-target markers OTUD5 and HK2, whose regulation highly correlated with the number of significantly downregulated proteins (Fig. 2b, Fig. S2). While **NE26380** and **NE26412** (Fig. 2b,c, Fig. S2) displayed IMiD-like degradation profiles against known neosubstrates, including ZFP91, CUL7 and ZBTB21, the remaining five analogues were almost inactive. Thus, the regioisomer SAR revealed a unique exit vector compatible with CDK4 degradation.

Maintaining the original regiochemistry of **NE25613**, we further probed the structure activity cliffs by a methyl scan and by single atom-modifications. These edits to the proximal segment substantially attenuated CDK4 degradation to less than 1.5-fold (entries 10-13), with one of the analogues (**NE26404**) specifically triggering a marked 6-fold decrease of RPRD1B levels instead. Minimal distal modifications (entries 14-17) also led to reduced CDK4 depletion while CRBN-independent off-target regulations where preserved. However, one structural analogue within this subset (**NE26396**) maintained robust activity, driving an ∼80% CDK4 depletion.

Complete removal of the distal shell in **NE26403** abrogated all cellular downregulations except for a very weak degradation effect on CDK4 (Fig. 2b,c). This residual activity was completely abolished when minimal modifications were introduced into the truncated structure (entries 19-25), further indicating that the proximal shell is an enabler of CDK4 recognition, while the distal region governs MGD potency, efficacy and selectivity. We therefore kept the proximal region and diversified the distal part with a variety of functional groups (entries 26-39). Among these analogues, we identified a major subset in which CDK4 degradation had been restored to more than 2-fold (entries 32-39). None of these analogues induced degradation of the alternative neosubstrate RPRD1B. All except two exhibited either no, or minimal CRBN-independent off-target activity, including **NE26381**, **NE26402** and **NE26394**, which elicited similar or even more pronounced CDK4 degradation than the primary screening hit **NE25613** (Fig. 2b-d, Fig. S2). Remarkably, **NE26381** simultaneously depleted CDK4, Schlafen 11 (SLFN11), a protein involved in immune and DNA damage responses,^42^ and the HIRA histone chaperone complex (HIRA, UBN1/2, CABIN1). MLN4924 co-treatment experiments and ubiquitinomics analysis mechanistically validated SLFN11 as a *bona fide* CRBN neosubstrate (Supplementary Fig. S3). This result exemplified how slightly different derivatizations can create entirely novel neosubstrate specificities, even in more advanced chemical optimization efforts. **NE26402** and **NE26394** exhibited striking selectivity toward CDK4, driving its efficient depletion by about 8-fold and 16-fold, respectively (Fig. 2b-d, Fig. S2). Apart from the dominating effect on CDK4, we detected very weak downregulations of the off-target TXNIP,^30^ the two known CDK4-binding partners INK4A and INK4C, and CDK6. Based on these assessments, we nominated **NE26394** as optimized hit compound for further evaluation.

We next confirmed that the depletion of CDK4 by **NE26394** was strictly dependent on the ubiquitin-proteasome system (UPS) and CRL4^CRBN^ activity, as demonstrated by co-treatment experiments with MLN4924 or the proteasome inhibitor bortezomib, as well as by evaluation of the compound in CRBN knockout cells (Supplementary Fig. S4). Moreover, compared to **NE25613**, the magnitude of **NE26394**-induced CDK4 ubiquitination was about two-fold higher (Fig. 3a, Supplementary Fig. S4). Concordantly, **NE26394** exhibited superior degradation potency and efficacy in parental HEK293 cells (DC_50_ = 163 nM, D_max_ = 93% vs. DC_50_ = 759 nM, D_max_ = 90% for **NE25613**), which were further amplified in CRBN-overexpressing cells (DC_50_ = 2.6 nM, D_max_ = 97%) (Fig. 2f, Fig. S4). The enhanced CDK4 degradation selectivity of **NE26394** also manifested as weaker downregulation of the close homologue CDK6 (DC_50_ ≫ 10 µM, observed D_max_ = 22%), representing at least a 60-fold preference for CDK4 over CDK6. In addition to CDK6, the kinases CDK1, CDK2, CDK5, CDK7, CDK9, GSK3α and GSK3β represent reported off-target liabilities of the CDK4 inhibitor atirmociclib, when administered at high concentrations.^12,13^ Notably, our lead molecule **NE26394** showed no activity against these kinases, further highlighting a potential selectivity advantage. In summary, these findings highlight the remarkable selectivity achievable with MGDs, maximizing CDK4 degradation while minimizing potential off-target liabilities.

**Fig. 3:**
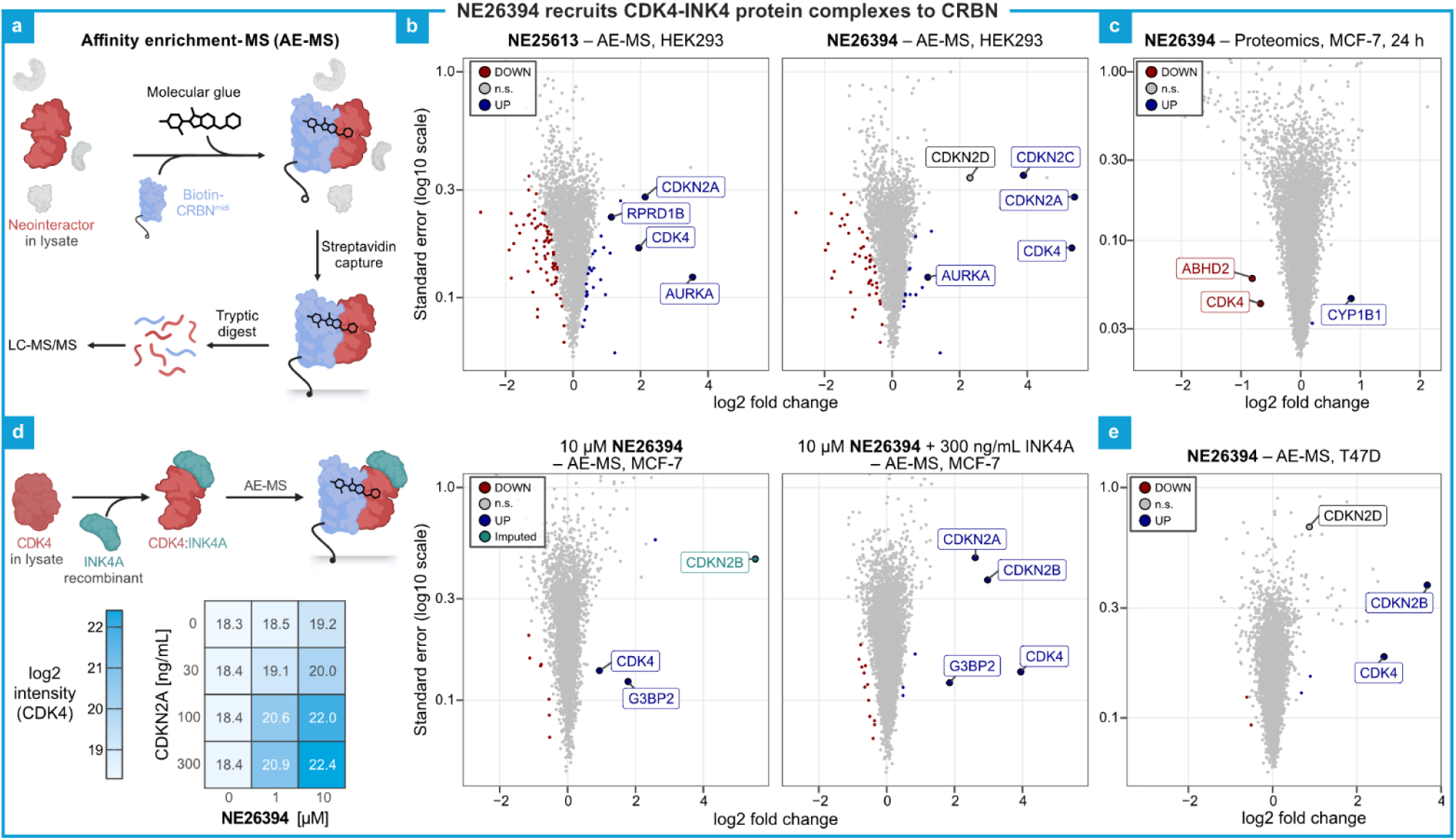
The analogue NE26394 is a selective CRBN-based molecular glue recruiting CDK4-INK4 complexes. **a**, Schematic for profiling compound-induced CRBN neointeractors using affinity-enrichment mass-spectrometry (AE-MS). **b**, AE-MS profiles of the screening hit **NE25613** (left) and the analogue **NE26394** (right) in HEK293 lysate demonstrating enrichment of INK4 proteins. **c**, Modest degradation of CDK4 by NE26394 in the breast cancer cell line MCF-7 (10 µM, 24 h). **d**, Spiking recombinant INK4A protein into MCF-7 lysate drastically enhances the molecular glue-induced CDK4 recruitment as observed by AE-MS. **e**, AE-MS profile of **NE26394** in the lysate of the breast cancer cell line T47D (10 µM). In volcano plots, log_2_ fold changes relative to vehicle (x-axis) and standard errors (y-axis, log_10_ scale) are shown. For AE-MS with **NE26394** and spiked-in INK4A, log_2_ fold changes relative to DMSO and INK4A spike-in are shown. Up- and downregulated proteins (mod. adj. p-value < 0.01) are coloured in blue and red, respectively. CDKN2B in the MCF-7 AE-MS experiment w/o recombinant INK4A was only detected in **NE26394**-treated samples and plotted with an imputed ratio coloured in turquoise. Not significant (n.s.) regulations are coloured grey.

### Involvement of INK4 family proteins in MGD-induced CDK4 recruitment to CRBN

Considering that neosubstrate ubiquitination by CRBN MGD provides strong evidence of induced cellular proximity to the CRL4^CRBN^ E3 ligase, the rapid ubiquitination kinetics on CDK4 are likely a direct consequence of ternary complexes formation promoted by our MGD compounds. To corroborate and characterize degrader-induced complexes involving CRBN and CDK4, we conducted in-lysate interactomics experiments employing affinity enrichment-mass spectrometry (AE-MS).^28–30^ Specifically, we incubated HEK293 cell extracts with our CDK4 degraders in the presence of biotinylated CRBN^midi^. CRBN^midi^ is a truncated, soluble CRBN construct optimized for biophysical assays that has also proven highly effective for AE-MS.^43^ Specific CRBN-MGD binders were subsequently captured via affinity purification and quantified by MS-based proteomics (Fig. 3a).

When assessing the screening hit **NE25613**, we recovered the two prominent neosubstrates CDK4 and RPRD1B (4- and 2.4-fold enrichment, respectively), alongside several other proteins (Fig. 3b). The most strongly enriched one among these interactors was the mitotic protein kinase Aurora kinase A (AURKA). This finding suggested the formation of a non-productive ternary complex, given that we did not detect any CRBN-dependent AURKA degradation in our cellular profiling experiments.^28^ The second most prominent interactor was the inhibitory CDK4-binding protein INK4A (CDKN2A), which was enriched to a degree comparable to CDK4. This finding points to the co-recruitment of INK4A into the CRBN-**NE25613**-CDK4 ternary complex.

Next, we tested **NE26394** and observed a remarkable, greater than 40-fold enrichment of CDK4, directly aligning with its enhanced cellular degradation efficacy (Fig. 3b). Conversely, RPRD1B and AURKA enrichment was almost completely abolished, confirming that our structural modifications successfully deselected these targets. **NE26394** promoted robust co-purification of the INK4A and INK4C (CDKN2C), which were also significantly downregulated in HEK293 cells (Fig. 2d). To a lesser extent, the related INK family member INK4D (CDKN2D) also was enriched, albeit without reaching statistical significance (Fig. 3b). A direct comparison of the AE-MS profiles for **NE25613** and **NE26394** revealed that INK4A co-recruitment closely correlated with the magnitude of CDK4 enrichment, suggesting the recruitment of pre-formed CDK4-INK4A heterodimers to CRBN. In contrast, other well-established CDK4 interactors, including D-type cyclins and the CIP/KIP family members p21, p27 and p57 were neither detected nor enriched in our AE-MS experiments. Thus, CDK4 complexes containing these regulatory subunits are evidently not recognized by CRBN. HEK293 cells were originally immortalized via transformation with human adenovirus 5 DNA, introducing the E1A oncoprotein to bind and sequester the RB tumour suppressor.^44^ Consequently, CDK4 activity is dispensable in HEK293 and does not control the expression of E2F target genes. We therefore shifted to therapeutically relevant, CDK4-dependent cell line models, such as the HR+ breast cancer cell line MCF-7, which is frequently used to evaluate the effect of CDK4 inhibition. Genetically, MCF-7 is characterized by deletions at the 9p21.3 gene locus, resulting in homozygous and hemizygous loss of CDKN2A and CDKN2B, respectively.^45^ Quantitative proteomics following a 24-hour **NE26394** treatment revealed a less than two-fold reduction of CDK4 in MCF-7 cells, in stark contrast to the near-complete CDK4 depletion in HEK293 cells. This suggested that low INK4 family protein abundance, particularly the lack of INK4A expression, severely reduced degradation efficacy. In line with this, AE-MS profiling using MCF-7 cell extracts and **NE26394** resulted in only a marginal, 1.8-fold enrichment of CDK4 with CRBN (Fig. 3d). We next sought to obtain functional validation for INK4-dependent CDK4 recruitment by supplementing MCF-7 lysates with increasing amounts of recombinant INK4A. Intriguingly, the addition of INK4A potentiated the interaction between CDK4 and CRBN in a dose-dependent manner, facilitating up to a 15-fold enrichment with concomitant purification of INK4A (Fig. 3d). The MCF-7 dataset further indicated that INK4B (CDKN2B), which was undetectable in HEK293 cells, can also acts as a CDK4 co-recruitment factor. We substantiated this finding via AE-MS using lysates from T47D cells, another widely used breast cancer cell line model known to be dependent on CDK4 activity.^12,46^ In contrast to the genetic deletions characterizing MCF-7, T47D cells lack INK4A expression due to epigenetic silencing of the CDKN2A gene, yet they retain expression of INK4B. AE-MS accordingly detected a much more prominent enrichment of CDK4 in T47D compared to MCF7 cells, an effect presumably mediated by INK4B (Fig. 3e). Thus, our AE-MS data provides substantial evidence that pairing of INK4 family proteins INK4A, INK4B or INK4C with CDK4 generates heterodimers that are specifically recognized as CRBN neosubstrates and represent the actual MGD target species in the cellular environment.

### A CDK4-selective MGD inhibits RB-E2F pathway activity in CDK4-dependent cancer cells

Compared to the weak CDK4 regulation in MCF-7, a 24-h treatment with **NE26394** in T47D cells resulted in an approximately 4-fold depletion of the kinase, consistent with enhanced CDK4 enrichment seen in our AE-MS experiments (Supplementary Fig. S5). Because CDK4-INK4B complexes are the major MGD substrates in these HR+ breast cancer cell lines, these results demonstrate that INK4B can compensate for the lack of INK4A to support degradation of CDK4. Next, we assessed the cellular effects of NE26394 in T47D cells. To enhance **NE26394**-mediated degradation potency and to minimize potential CRBN/CRL-unrelated off-target activity, we moderately overexpressed CRBN in these cells (∼8-fold increase). Treatment of T47D-CRBN^oe^ cells with 1 µM **NE26394** resulted in a 4-fold CDK4 downregulation, mirroring the degradation observed in parental T47D cells at a ten-fold higher compound concentration.

To compare RB-E2F pathway perturbations elicited by either **NE26394** or the CDK4 inhibitor atirmociclib, we performed 24-h treatments with both compounds. Intriguingly, except for the specific downregulation of CDK4 and INK4B (CDKN2B) in **NE26394**-treated cells, the recorded proteomic profiles for both compounds were almost identical. Both datasets revealed a comparable decrease in the abundance of key E2F target proteins, including PLK1, RRM2, TK1 and cyclin A2 (CCNA2) (Fig. 4a).

**Fig. 4:**
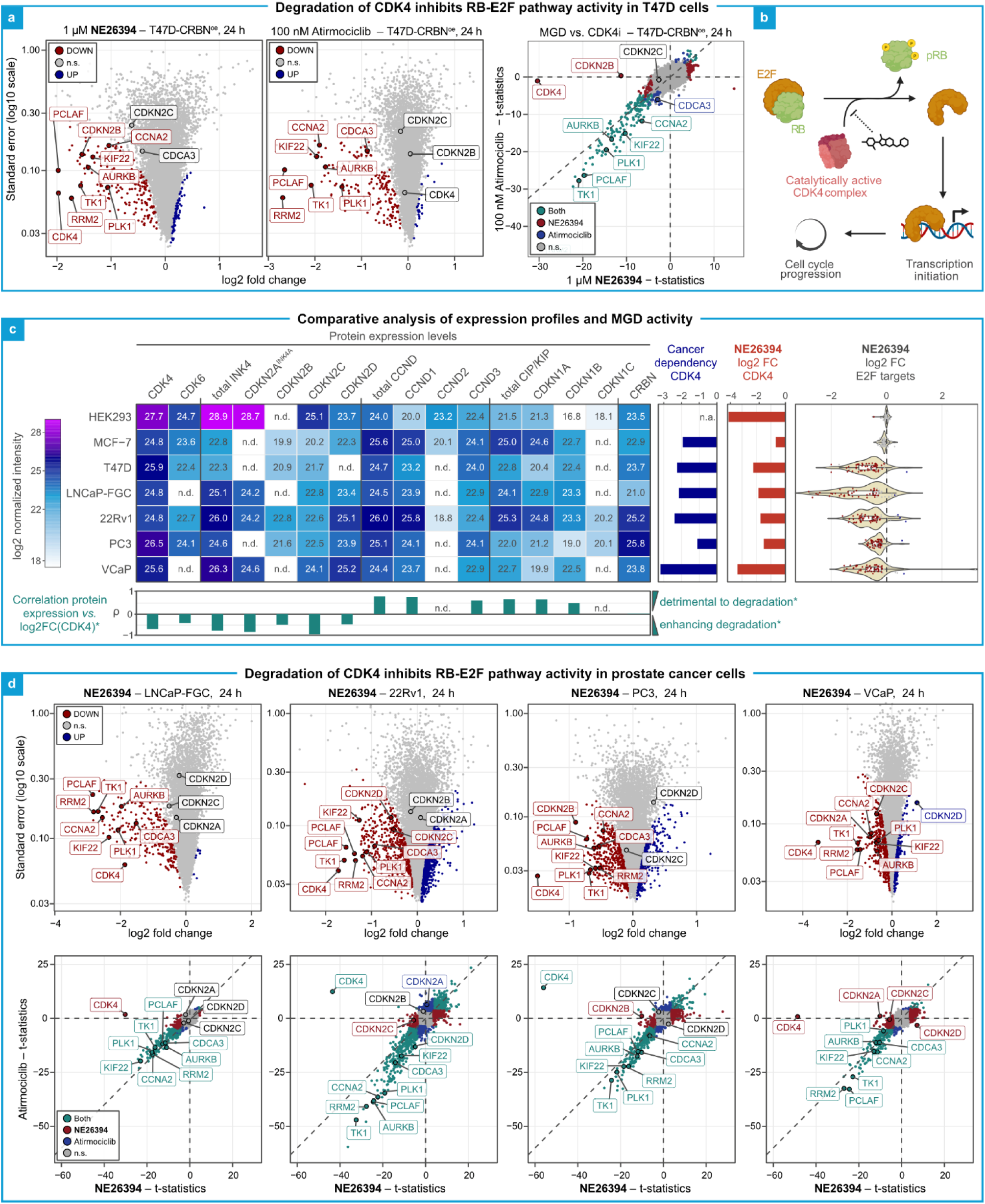
CDK4 MGD NE26394 inhibits RB-E2F pathway activity in disease-relevant cancer cell lines. **a**, Proteome-wide regulations following 24 h treatment of T47D-CRBN^oe^ cells with 1 µM **NE26394** (left) or 100 nM atirmociclib (centre), and comparison of their t-statistics (right). **b**, Scheme for CDK4-Rb-E2F pathway. **c**, Expression levels of various CDK4-related proteins and protein families across seven cell lines (heatmap). Horizontal bar charts display the dependency of those cell lines on CDK4 for survival and growth (DepMap CRISPR score^46^; left) and the extent of downregulation induced in them by treatment with 10 µM **NE26394** for 24 h (log_2_ fold change; right). The treatment-associated changes in protein levels of validated target genes of E2F-induced transcription^47^ are depicted as violin plots (log_2_ fold changes). Pearson correlations of protein expression levels with compound-induced depletion of CDK4 are represented below the heatmap; A negative correlation coefficient ρ suggests high protein expression associates with strong CDK4 degradation, whereas ρ ≈ 0 and ρ → 1 correspond to no and detrimental effects, respectively. **\***These correlations should be interpreted cautiously, as the small size of the cell line panel limits statistical robustness. **d**, Proteomic regulations following 24 h treatment of cell culture models of prostate cancer with 10 µM **NE26394** (top) and comparison to those induced by 100 nM CDK4i atirmociclib (bottom). In the volcano plots, log_2_ fold changes relative to vehicle (x-axis) and standard errors (y-axis, log_10_ scale) are shown. Up-and downregulated proteins (mod. adj. p-value < 0.01) are coloured in blue and red, respectively. Non-significant (n.s.) regulations are coloured in grey. In the comparison plot, the t-statistics for regulations in two different proteomics experiments are plotted against each other. Statistically significant up- and downregulations in the x- and y-axis experiments are coloured in blue and red, respectively, overlapping significant features are coloured turquoise, and non-significant regulations are coloured grey.

Besides HR+ breast cancer, prostate adenocarcinoma has been reported to heavily rely on CDK4 for survival, with limited expression and a non-essential role of CDK6.^12,46^ We therefore assessed proteomic expression profiles of four CDK4-dependent and RB-proficient prostate cancer cell lines, which were of different origins and androgen receptor status (LNCaP-FGC, 22Rv1, PC3 and VCaP). Quantitative comparisons with HEK293, MCF-7 and T47D proteomes revealed highest INK4 protein expression in HEK293 cells. While the abundance of individual INK4 proteins varied significantly across the prostate cancer cell lines, overall levels were considerably higher than in the breast cancer cell lines MCF-7 and T47D (Fig. 4b). Thus, we reasoned that the selected prostate cancer cell lines should respond well to **NE26394** in terms of CDK4 degradation and RB-E2F pathway modulation. We detected robust CDK4 depletion across these models, ranging from a 3-fold reduction in PC3 to a 10-fold depletion in VCaP cells (Fig 4b,c). **NE26394** treatment broadly suppressed E2F target protein expression across all four cell lines, with the most pronounced downregulation seen in LNCaP-FGC cells (Fig 4b,c). Consistently, quantitative proteomics uncovered highly correlated proteomic profiles when comparing 24-h treatments with either our MGD compound **NE26394** or the kinase inhibitor atirmociclib. These data underscore that our degrader compound effectively suppresses therapeutically relevant, CDK4-mediated signalling across diverse prostate cancer cells.

Finally, to get first insights into potential markers of **NE26394** sensitivity, we explored correlations between the extent of CDK4 degradation and the expression of CDK4 binding partners based on the proteome quantifications across the seven cell lines used in this study (Fig. 4c, Supplementary Fig. S6). Reassuringly, high INK4 protein levels, particularly of INK4A (CDKN2A) and INK4C (CDKN2C), appeared to correlate well with strong CDK4 degradation. The available data further indicated an inverse correlation for D-type cyclin expression, which may reflect competition with INK4 proteins for cellular CDK4 binding. A similar, somewhat weaker trend emerged for p21 (CDKN1A) and p27 (CDKN1B), possibly due to their stabilizing influence on cyclin D-CDK4 complexes.^48^ While a conclusive analysis will require a larger cell line panel, these initial findings outline a path toward identifying the mechanistic determinants of CDK4 degradation, while laying the groundwork to evaluate its impact on RB-E2F pathway activity and cell cycle progression.

## Discussion

We report the first CRBN-directed MGD degraders of the cancer target CDK4, supported by comprehensive evidence of specific, MGD-induced recruitment of CDK4 into CRBN-MGD complexes *in vitro* and rapid MGD-promoted ubiquitination and CRBN/CRL-dependent CDK4 degradation in intact cells. Starting from a singleton hit in a target-agnostic proteomics screening campaign of several thousand CRBN MGD compounds, we generated a series of analogues and systematically derived SAR data based on their disparate proteomic profiles with CRL4^CRBN^-dependent and -independent off-target regulations. Through this, we identified **NE26394** that degraded CDK4 selectively, more potently and with higher efficacy than our screening hit **NE25613**, owing to stronger ternary complex formation and ubiquitination.

Our AE-MS interactomics experiments using **NE26394** and recombinant CRBN bait demonstrated co-enrichment of CDK4 binding partners from the inhibitory INK4 family proteins, such as the tumour suppressor protein INK4A. Remarkably, as neither activating D-type cyclins nor CIP/KIP family CDK inhibitory proteins (*e.g.*, p21, p27) were affinity-purified by **NE26394**, our results pointed to CDK4-INK4A and CDK4-INK4C heterodimers as specific neosubstrate species in the HEK293 cellular background. However, this finding did not exclude specific recruitment of CDK4 monomers into ternary complexes with CRBN and CDK4-directed MGDs. While CDK4 was only weakly enriched in AE-MS experiments with lysate from INK4A-negative MCF-7 cells,^45^ spiking in recombinant INK4A led to a strong and dose-dependent increase in MGD-induced CDK4 binding. Thus, these findings argue against the direct recognition of CDK4 monomers and instead may suggest an INK4-induced allosteric rearrangement upon CDK4 binding, promoting surface complementarity and heterodimer recruitment into quaternary CRBN-MGD-CDK4-INK4 protein complexes.^49^ While co-recruitment and co-depletion of direct CRBN neosubstrates and their interaction partners (referred to as “bystander” neosubstrates) have been reported previously,^26,28,50^ structural data have demonstrated that CRBN-MGD engagement of neosubstrates like CSNK1A1 or G3BP2 occurs independently of their associated “bystanders”, FAM83G and USP10, respectively. In contrast, INK4 proteins appear to act as essential auxiliary subunits in heterodimeric complexes with CDK4. They may therefore be indispensable for CDK4 recruitment and subsequent degradation, constituting a novel mechanism governing specific CRBN neosubstrate engagement. Further experimental work will be needed to explore the underlying structural determinants – in particular, whether the CRBN-MGD binding interface is created *de novo* upon INK4 protein-induced conformational rearrangement of the CDK4 protein surface, or whether a composite contact site involving both CDK4 and INK4 protein surface elements may be formed.

Association of different INK4 proteins with CDK4 or CDK6 is thought to occur with comparable affinity and trigger similar structural changes, which interfere with CDK4/6 kinase activity by preventing both ATP and D-type cyclin binding.^49^ However, even if this applies under defined conditions *in vitro*, the impact of individual INK4 family members on CDK4 degradation may vary across different cellular environments. Functional outcomes may depend on INK4 protein levels and on various post-translational modification, competitive binding of other CDK4-interacting factors like cyclin D1, or additional regulation events.^7,51,52^ Our data suggests a critical role of INK4A, INK4B and INK4C for robust, MGD-induced CDK4 degradation, as we found them affinity-purified and degraded together with CDK4 in AE-MS and global proteomics, respectively. Intriguingly, p16^INK4A^ represents a rare example of a lysine-free protein, meaning it cannot serve as direct substrate of CRL4^CRBN^ E3 ligase-mediated ubiquitination via conventional lysine modification.^53^ We hypothesize that the observed weak downregulation of p16^INK4A^ following MGD exposure reflects secondary destabilization of monomers liberated upon cellular CDK4 depletion, possibly due to enhanced alternative N-terminal ubiquitination by a constitutive cellular E3 ligase other than CRL4^CRBN^.^53^ On the other hand, INK4B and INK4C do harbour lysines that are ubiquitinated upon MGD treatment (data not shown), qualifying them as associated ubiquitination targets of functional CRBN ternary complexes. Compared to CDK4, the INK4 proteins are degraded to a much lesser extent. Thus, INK4 proteins released from CDK4 may then either sequester monomeric CDK4 or compete with D-type cyclins to induce the dissociation of e.g. CDK4-cyclin D1 complexes, resulting in the recursive formation of CDK4-INK4 protein dimers serving as CRBN-MGD neosubstrates in subsequent rounds of degradation.^54^ According to such a model, the minimal reduction seen for INK4A in MGD treatment experiments may potentially trigger deeper CDK4 degradation than supported by INK4B, for which we detected a decrease in protein amounts by about 50% in T47D or PC3 cells. However, as even INK4B is moderately regulated in comparison to MGD-induced CDK4 depletion, INK4 proteins may be sustained at sufficiently high levels to efficiently co-catalyse the MGD-promoted CDK4 degradation. Importantly, relative to CDK4 expression, such a model implies that only sub-stoichiometric INK4 protein amounts are needed, a criterion that may even be met in tumours harbouring CDKN2A and CDKN2B deletions, considering the functional redundancy among INK4 family members.

Our optimized hit compound **NE26394** induced proteomic changes associated with E2F-mediated pathway inhibition, similar to the next-generation CDK4 inhibitor atirmociclib in therapeutically relevant, CDK4-dependent cell line models. Although steep activity cliffs can complicate optimizing MGD on-target and minimizing their off-target effects, they also add an additional layer of selectivity compared to ATP-competitive inhibitors, which often struggle to differentiate the largely conserved catalytic pocket. In contrast, **NE26394** displays remarkable selectivity over other related protein kinases, including the close homologue CDK6 and CDK family members that are inhibited by higher doses of atirmociclib.^12^ Moreover, due to prolonged target inactivation that outlasts the drug’s presence, the event-driven pharmacology of degrader molecules has the potential to translate into profound pharmacokinetic advantages over “occupancy-driven” small molecule inhibitors. While our novel CDK4 MGDs are at an early stage, we have demonstrated they are amenable to medicinal chemistry optimization, resulting in the identification of drug-like, selective CDK4 degraders compliant with Lipinski’s rule of five.^55^ We expect that their degradation potency and efficacy can be further improved to exploit their full potential as therapeutic agents.

## Methods

### Reagents

MLN4924, bortezomib, 2-chloroacetamide (CAA), tris(2-carboxyethyl)phosphine hydrochloride (TCEP), sodium deoxycholate, Na_2_HPO_4_, 3-(N-morpholino)propanesulfonic acid (MOPS), Tris(hydroxymethyl)-aminomethan, NP-40 alternative, glycerol, trifluoracetic acid, sodium chloride, formic acid and acetonitrile were from Merck. n-Undecyl-β-D-maltoside (UDM) was from Anatrace. Protease inhibitor mix was from ThermoFisher Scientific (A32955). Trypsin was from Promega. PTMScan^®^ HS Ubiquitin/SUMO Remnant Motif (K-ε-GG) Kit (#59322) was from Cell Signaling Technology. Atirmociclib, palbociclib and recombinant INK4A (HY-P72785) were from MedChemExpress.

### Cell culture, drug treatments and cell lysis (Proteomics)

All cell lines were from Cytion. HEK293 cells, their derivatives (CRBN knockout and CRBN overexpression), and MCF-7 were cultured in DMEM (VWR) supplemented with 10% FCS (Thermo Fisher Scientific). VCaP were cultured in DMEM high glucose (VWR) with 10% FCS. T47D, 22RV1, LNCaP-FGC were cultured in RPMI-1640 (VWR) supplemented with 10% FCS. PC3 were cultured in 1:1 DMEM+F12 (VWR) with 5% FCS. All compounds were dissolved in DMSO, to generate 1000× stock solutions. Compounds were plated in 96-well plates according to a randomized layout using an Opentrons OT-2. 40 (screening) or 27 (for most follow up experiments) compounds were allocated on each plate (in duplicate or triplicate, respectively), along with 16 or 15 DMSO controls, respectively. Cells were treated at 10 µM (unless specified otherwise) for the specified amount of time, washed with PBS and harvested using NEOsphere lysis buffer (0.05% UDM, 75 mM Tris-HCl pH 8.5, 40 mM CAA, 10 mM TCEP). The lysates were heated to 80 °C for 10 min while shaking in a Thermomixer (Eppendorf). Proteins were digested by adding 400 ng of trypsin per sample (overnight, 37 °C). The resulting peptides were desalted using in-house prepared, 200 µL two plug C18 StageTips (3M Empore^TM^)^56^ and then analysed by LC-MS/MS.

### Generation of stable cell lines

The plasmid vectors for stable cell line generation (overexpression of CRBN (CRBN^oe^) and CRBN knockout (CRBN^KO^)) were from VectorBuilder. The guide RNA sequence for establishing CRBN-/-cells was as follows: ATCTAACTTCATGGCCTCGCGTTTTAGAGCTAGAAATAGCAAGTTAAAA-TAAGGCTAGTC CGTTATCAACTTGAAAAAGTGGCACCGAGTCGGTGC. HEK293 cells with stable gene overexpression or CRBN knockout were generated by lentiviral-mediated infection of cells according to published protocols.^57^ In brief, HEK293T were co-transfected with psPAX2, pMD2.G, pRSV-Rev and a lentiviral construct for CRBN expression (CRBN^oe^) or Cas9 and gRNA (CRBN^KO^) under the control of a EF1A (CRBN_OE) or U6 (CRBN_KO) promoter. The supernatant was harvested after 48 h of cell transfection and passed through a 0.45 µm filter. HEK293 cells were infected with the supernatant for 24 h, followed by selection of transduced cells with puromycin (2 µg/mL). CRBN overexpression and knockout were verified by MS-based proteome profiling.

### Ubiquitinomics

Cell treatment and sample preparation was done according to our recently established protocol with a few modifications.^38^ In brief, cells were cultured in 12-well plates in respective media and treated for 30 min with the specified compounds, followed by lysis with SDC buffer. Protein concentrations were determined using the BCA assay (Merck-Millipore) and the proteins were digested overnight at 37°C using 100:1 protein:trypsin ratio (Promega). After digestion, immunoprecipitation (IP) buffer (50 mM MOPS pH 7.2, 10 mM Na_2_HPO_4_, 50 mM NaCl) was added to the samples together with K-GG antibody-bead conjugate, followed by a 2 h incubation on a rotor wheel. Beads washing and peptide elution was performed according to manufacturer’s instructions. The peptide eluate was desalted using in-house prepared, 200 µL two plug C_18_ StageTips (3M EMPORE^TM^).^56^

### Affinity enrichment-mass spectrometry

Cells were lysed in ice-cold NP-40 buffer (0.05% NP-40, 50 mM Tris-HCl pH 7.5, 150 mM NaCl, 5% glycerol), freshly supplemented with protease inhibitors, followed by sonication and centrifugation at 20,000 ×g for 10 min (4 °C). The supernatant was transferred to a fresh tube. The protein concentration was determined using a BCA assay kit (Merck-Millipore) and the lysate concentration was adjusted to 1 mg/mL. Biotinylated CRBN-midi (CRELUX, WuXi AppTec) was added to the lysate along with DMSO/compound and incubated for 1 h at 4 °C. Biotin affinity capture was used to isolate CRBN-glue-bound proteins (4 washes with lysis buffer after enrichment). Proteins were eluted with NEOsphere lysis buffer and digested by adding 100 ng of trypsin per sample (overnight, 37 °C). The resulting peptides were desalted using in-house prepared, 200 µL two plug C18 StageTips (3M Empore^TM^)^56^ and then analysed by LC-MS/MS.

### LC-MS/MS measurements

Peptides were either analysed on mass spectrometers from Bruker (timsTOF pro2, timsTOF HT or timsTOF Ultra 2/timsAIP) or ThermoFisher (Orbitrap Astral or Orbitrap Astral Zoom). The LC Setup differed for the various sample types and/or mass spectrometers. For global proteomics on timsTOF instruments and global ubiquitinomics the following LC setup was used: Peptides were loaded on 30 cm reverse-phase columns (75 µm inner diameter, packed inhouse with ReproSil Saphir 100 C18 1.5 µm resin [Dr. Maisch GmbH]) using either a Vanquish™ Neo system (ThermoFisher) or a nanoElute® 2 system (Bruker). The column temperature was maintained at 60°C using a column oven. The LC flow rate was 300 nL/min and the complete gradient was 50 min (global proteomics) or 45 min (ubiquitinomics). The LC setup of the samples measured with a higher throughput (global proteomics samples measured on the Orbitrap Astral (or Zoom) instrument at ∼65 samples per day (SPD) or AE-MS samples measured at ∼100 SPD differed as follows: a 17 cm column with a 150 µm inner diameter was used, the flow rate was at 2,000 nL/min and the gradient length was 17 min (global proteomics) or 9 min (AE-MS), respectively. Data acquisition on timsTOF instruments was done using diaPASEF^58^ (proteomics) or slicePASEF^59^ (for ubiquitinomics). The diaPASEF acquisition scheme (ion mobility (IM) range from 0.65-1.35 and m/z range from 300-1,500) and was generated with py_diAID^60^. For global proteomics data measured on the Orbitrap Astral, a Nanospray Flex™ source was used, with an ionization voltage of +2.2 kV, and an ion transfer tube temperature of 280°C. The Orbitrap scan resolution was 240,000. The RF lens value was 40%, the AGC target 500%, and the maximum injection time 3 ms. A data-independent acquisition (DIA) scheme with 222 m/z windows of 3 to 10 Th covering an m/z range of 320-1231 was used, with the following Astral settings: scan range: 150 - 2,000 m/z, normalized HCD collision energy: 25%, AGC target: 800%, RF lens: 40%, maximum injection time: 3 ms.

### Raw data processing

MS raw files were analysed using DIA-NN^61^ v2.1 enterprise (proteomics) or 2.5 enterprise (ubiquitinomics and AE-MS). Reviewed UniProt entries (human, SwissProt 01-2023 [9606]) were used as protein sequence database for DIA-NN searches. For proteomics and interactomics (AE-MS), one missed cleavage, a maximum of one variable modification (oxidation of methionines) and N-terminal excision of methionine were allowed. For ubiquitinomics, the number of missed cleavages and the maximum number of variable modifications was set to 2. K-GG (UniMod:121) and N-terminal acetylation (UniMod:1) were configured as variable modifications. Carbamidomethylation of cysteines (UniMod:4) was set as fixed modification for all analyses. All data processings were carried out using library-free analysis mode in DIA-NN. --tims-scan was added as an additional command in case of ubiquitinomics.

### Statistical data analysis

DIA-NN outputs were further processed with R. Complete missing cases in any of the conditions tested were rescued by accepting low-quality precursors (i.e., q-value > 0.01), where possible. For ubiquitinomics, missing values were imputed in DMSO in case of complete missingness. To minimize false positives, imputation was performed only when 100% condition completeness was achieved for a given peptide in at least one compound-treated condition. K-GG peptide to site mapping was done using reviewed entries of the human UniProt database (SwissProt, release 01-2023). The protein (or peptide) intensities were normalized by median scaling and corrected for variance drift over time (if present) using the principal components (derived from principal component analysis (PCA)) belonging to DMSO samples. Subsequently, protein (or peptide, for ubiquitinomics) intensities were subjected to statistical testing with variance and log fold-change moderation using LIMMA.^62^ p-values corrected for multiple hypothesis testing were used to assess significance for proteomics (adjusted p-value < 0.01) and ubiquitinomics (adjusted p-value < 0.05). For comparing proteome and ubiquitinome data, identifications were mapped at the protein level.

Protein intensity levels across cell lines were obtained from DIA-NN label-free quantification (separate search runs). For gene families (INK4, D-type cyclins, and CIP/KIP), family-level intensity values were calculated by summing the intensities of detected members. Associations between protein abundance and CDK4 fold-change upon **NE26394** treatment were assessed by calculating Pearson correlations between each protein’s median-centred, log_2_-transformed intensity and CDK4 log_2_ fold-change following **NE26394** treatment across cell lines. Distributions of E2F target gene fold changes following **NE26394** treatment were also compared across cell lines.

## Data availability

DIA-NN is freely available for academic use and can be downloaded at https://www.github.com/vdemichev/diann.

Cancer dependency and CRISPR screening data were sourced from the DepMap portal (DepMap Public 26Q1) at https://depmap.org/.

## Acknowledgements

The authors gratefully acknowledge Dr. Lieven Meerpoel for valuable discussions regarding the medicinal chemistry design.

Figures 1a (YO29UYE25T), 3a (WY29UYED3R), 3d (OB29UYE8PT), 4b (HO29UYEIG7) were generated with BioRender (Agreement number RM27FBDPFP).

## Author contributions

Conceptualization and experiment design: P.R.A.Z., H.D., M.S.

Development of data analysis pipeline: B.Sch., I.S., B.Sh.

Data analysis and interpretation: P.R.A.Z., H.D., M.S., B.Sh.

Figure preparation: P.R.A.Z., B.Sh., M.S.

Manuscript writing: P.R.A.Z., H.D.

Proofreading and editing of the Manuscript: B.Sh., M.S., U.O., I.S.

Execution of experiments: A.B., D.B., S.M., D.W., T.G.

All authors read and approved the final manuscript.

## Author information

Correspondence and requests for materials should be addressed to

## Supplementary Figures

**Fig. S1:**
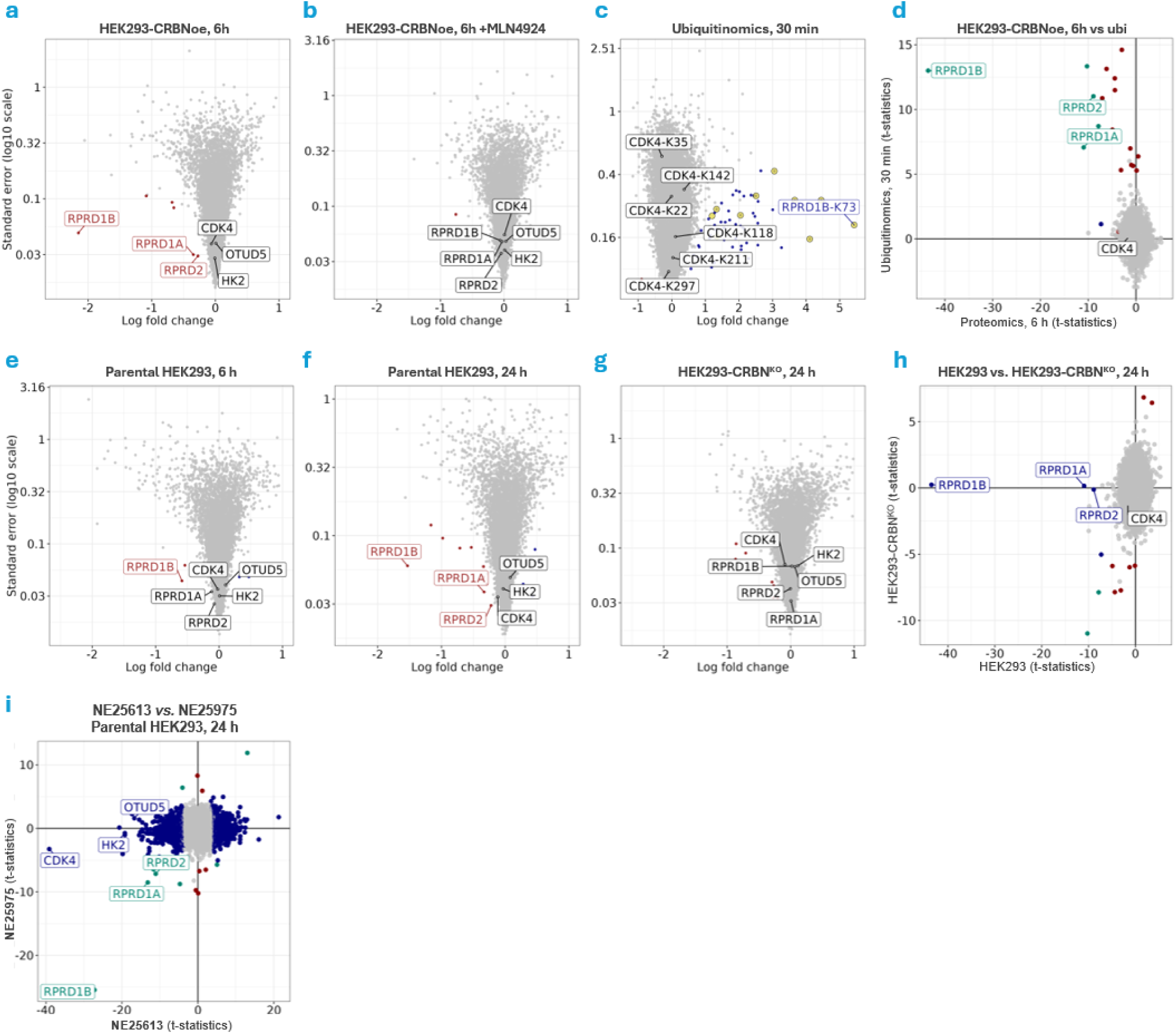
Mechanistic validation of the additional MGD primary screening hit NE25975 targeting RPRD1B. **a-d**, Compound-induced regulations in HEK293-CRBN^oe^ cells. Proteomic regulations in HEK293-CRBN^oe^ induced by **NE25975** alone (**a**) or in co-treatment with the neddylation inhibitor MLN4924 (**b**) after 6 h. Induced ubiquitination of RPRD1B and associated proteins RPRD1A and RPRD2 after 30 min (**c**) and t-statistic comparison of global proteomics and ubiquitinomics regulations (**d**), highlighting RPRD1B as the most prominently ubiquitinated and degraded neosubstrate. **e,f** Regulations in parental HEK293 cells after 6 h (**e**) and 24 h (**f**). **g,h**, Regulations in HEK293-CRBN^KO^ cells after 24 h (**g**) and t-statistic comparison with parental HEK293 cells (**h**), highlighting CRBN-dependent depletion of RPRD proteins as well as CRBN-independent regulations. **i**, t-statistic comparison of protein regulations of primary screening hits **NE25613** and **NE25975** in HEK293 cells after 24 h. While both MGDs degrade RPRD1B to similar extent, **NE25975** does not degrade CDK4 or exhibit the broader CRL4^CRBN^-independent proteome perturbations. In volcano plots, log_2_ fold changes relative to vehicle (x-axis) and standard errors (y-axis, log_10_ scale) are shown. For MLN4924 co-treatment, fold-changes relative to vehicle + MLN4924 are shown. Up- and downregulated proteins (mod. adj. p-value < 0.01) or ubiquitination sites (mod. adj. p-value < 0.05) are coloured in blue and red, respectively. Non-significant (n.s.) regulations are coloured grey. In comparison plots, t-statistics for regulations in two different experiments are plotted against each other. For comparison of global proteomics and ubiquitinomics profiles, ubiquitination sites mapping to the same protein were averaged, and only significantly upregulated sites were included in the analysis. Statistically significant up-and downregulations in the x- and y-axis experiments are coloured in blue and red, respectively, overlapping significant features are coloured turquoise, and not significant regulations are coloured grey.

**Fig. S2:**
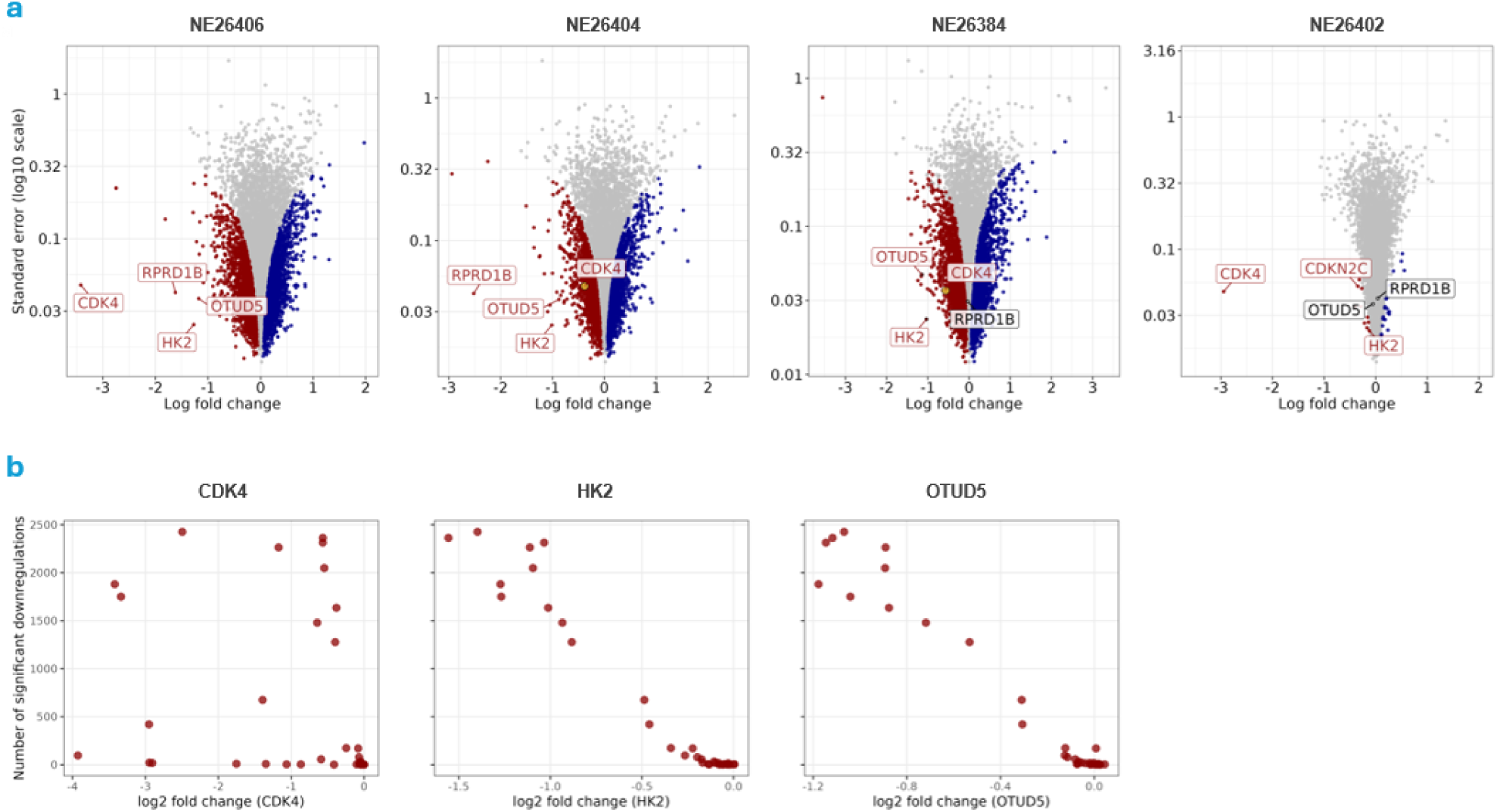
Further analysis of SAR datasets in HEK293 cells following 24 h treatment with MGD analogues. **a**, Additional proteomic profiles of selected analogues: Hit enantiomer **NE26406** mirrors regulations of the racemate and its enantiomer **NE26408**. Minimal edits to the proximal sphere in **NE26404** resulted in favoured RPRD1B degradation. Substitution in the distal sphere abrogated CDK4 and RPRD1B degradation for **NE26384** but maintained CRBN-independent regulations, whereas for **NE26402**, it resulted in potent and selective CDK4 degradation. **b**, Correlation of the indicated protein regulations with the total number of downregulations for individual compounds. In the volcano plots, log_2_ fold changes relative to vehicle (x-axis) and standard errors (y-axis, log_10_ scale) are shown. Up-and downregulated proteins (mod. adj. p-value < 0.01) or ubiquitination sites (mod. adj. p-value < 0.05) are coloured in blue and red, respectively. Not significant (n.s.) regulations are coloured grey.

**Fig. S3:**
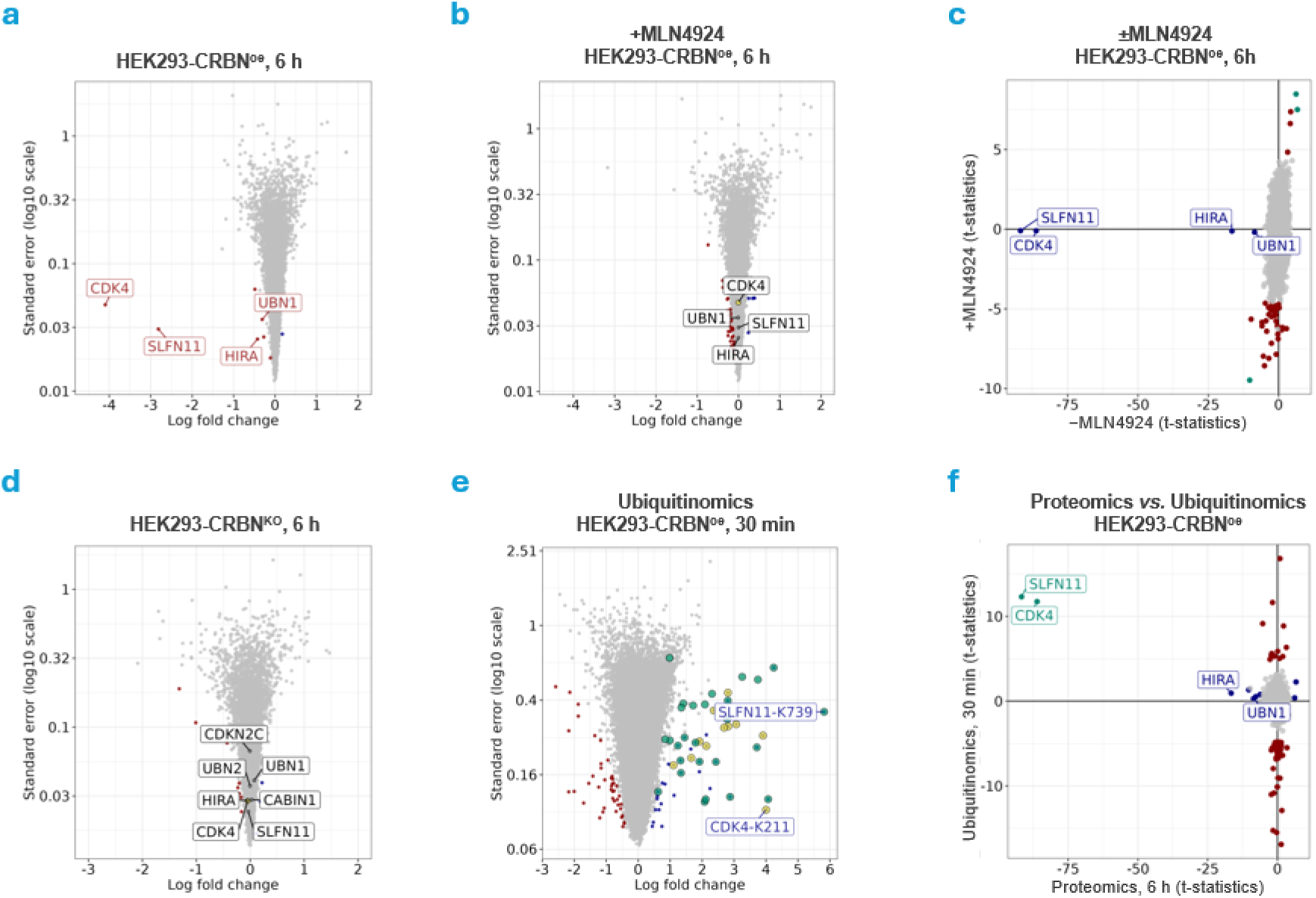
Mechanistic validation of SLFN11 MGD NE26381 in HEK293 cell lines. **a**,**b**, Proteomic regulations in HEK293-CRBN^oe^ induced by **NE26381** alone (**a**) or in co-treatment with the neddylation inhibitor MLN4924 (**b**) after 6 h. **c**, t-statistic comparison of ±MLN4924 treatments. **d**, Loss of SLFN11 and CDK4 regulations in HEK293-CRBN^KO^ cells. **e**,**f** Induced ubiquitination of SLFN11 (highlighted green) and CDK4 (highlighted yellow) after 30 min (**e**) and t-statistic comparison of global proteomics and ubiquitinomics regulations (**f**), highlighting SLFN11 and CDK4 as the most prominently ubiquitinated and degraded neosubstrates. In the volcano plots, log_2_ fold changes relative to vehicle (x-axis) and standard errors (y-axis, log_10_ scale) are shown. For MLN4924 co-treatment, fold-changes relative to vehicle + the respective inhibitor are shown. Up- and downregulated proteins (mod. adj. p-value < 0.01) or ubiquitination sites (mod. adj. p-value < 0.05) are coloured in blue and red, respectively. Non-significant (n.s.) regulations are coloured grey. In comparison plots, t-statistics for regulations in two different experiments are plotted against each other. For comparison of global proteomics and ubiquitinomics profiles, ubiquitination sites mapping to the same protein were averaged, and only significantly upregulated sites were included in the analysis. Statistically significant up-and downregulations in the x- and y-axis experiments are coloured in blue and red, respectively, overlapping significant features are coloured turquoise, and non-significant regulations are coloured grey.

**Fig. S4:**
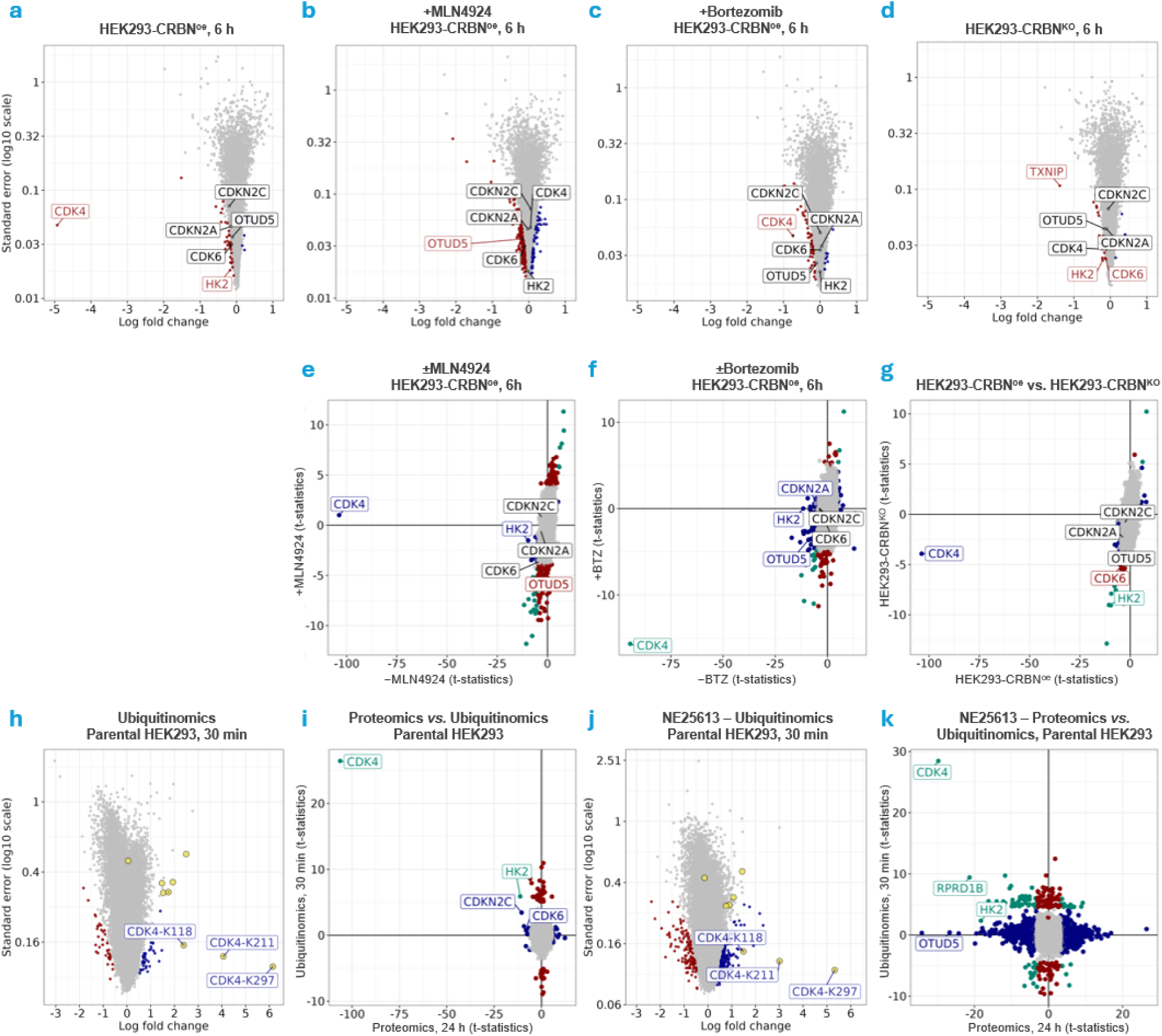
Mechanistic validation of CDK4 MGD NE26394. **a-c**, Volcano plot visualizations (log_2_ fold-change (x-axis) versus log_10_ standard error (y-axis) of the proteome regulations of HEK293-CRBN^oe^ cells treated for 6 h with **NE26394** alone (**a**), or co-treated with MLN4924 (**b**), or bortezomib (**c**). **d**, Proteomic regulations following 6 h treatment of HEK293-CRBN^KO^ cells with **NE26394**. **e-g**, t-statistic comparisons of proteomics analyses of **NE26394** ±MLN4924 (**e**) or ±bortezomib (**f**) in HEK293-CRBN^oe^ cells, as well as comparison of **NE26394** regulations in HEK293-CRBN^oe^ and HEK293-CRBN^KO^ cells (**g**). **h,j**, Induced CDK4 ubiquitination sites (highlighted in yellow) in parental HEK293 following treatment with **NE26394** and **NE25613**, respectively. **i,k**, t-statistic comparisons of proteomic regulations after 24 h, and ubiquitinomics regulations after 30 min treatment of parental HEK293 following treatment with **NE26394** and **NE25613**, respectively. In thr volcano plots, log_2_ fold changes relative to vehicle (x-axis) and standard errors (y-axis, log_10_ scale) are shown. For MLN4924 or bortezomib co-treatments, fold-changes relative to vehicle + the respective inhibitor are shown. Up- and downregulated proteins (mod. adj. p-value < 0.01) or ubiquitination sites (mod. adj. p-value < 0.05) are coloured in blue and red, respectively. Non-significant (n.s.) regulations are coloured grey. In comparison plots, t-statistics for regulations in two different experiments are plotted against each other. For comparison of global proteomics and ubiquitinomics profiles, ubiquitination sites mapping to the same protein were averaged, and only significantly upregulated sites were included in the analysis. Statistically significant up-and downregulations in the x- and y-axis experiments are coloured in blue and red, respectively, overlapping significant features are coloured turquoise, and non-significant regulations are coloured grey.

**Fig. S5:**
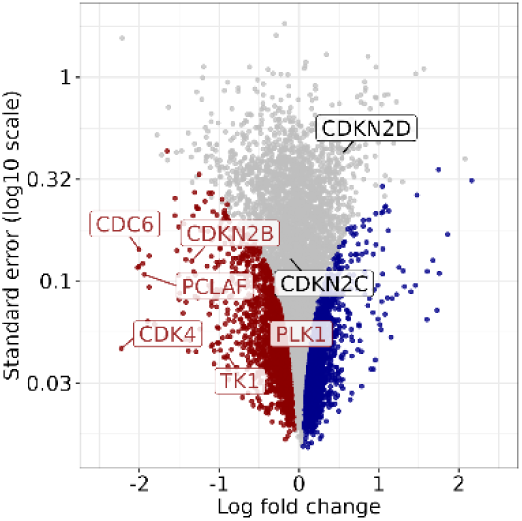
Proteomic regulations in parental T47D cells after treatment with 10 µM NE26394 for 24 h. log_2_ fold changes relative to vehicle (x-axis) and standard errors (y-axis, log_10_ scale) are shown. Up- and downregulated proteins (mod. adj. p-value < 0.01) are coloured in blue and red, respectively. Not significant (n.s.) regulations are coloured grey.

**Fig. S6:**
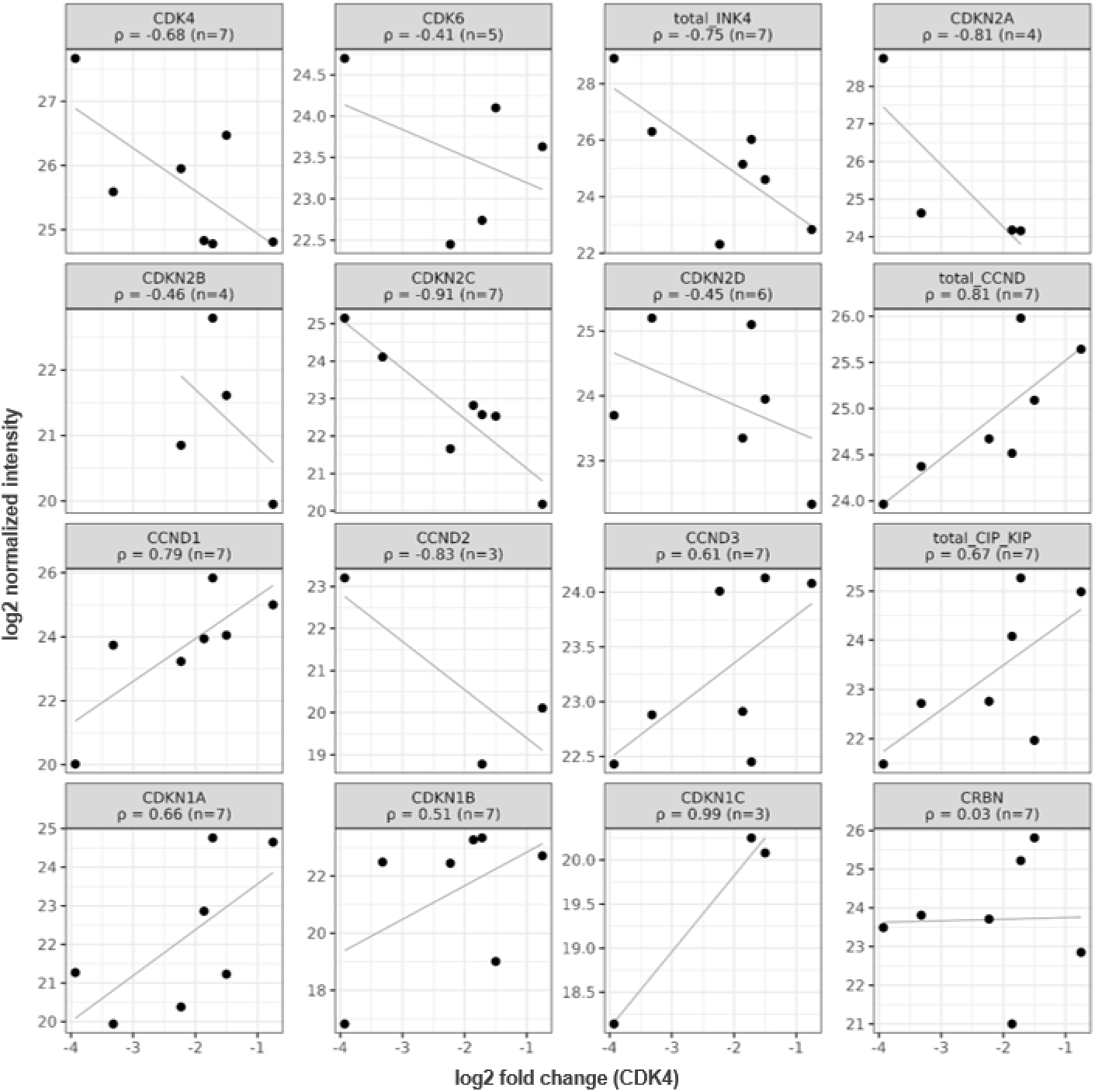
Pearson correlations of selected protein expression levels with NE26394-mediated CDK4 depletion.

## References

1. Sherr, C. J., Beach, D. & Shapiro, G. I. Targeting CDK4 and CDK6: From Discovery to Therapy. Cancer Discov. 6, 353–367 (2016).

2. Fassl, A., Geng, Y. & Sicinski, P. CDK4 and CDK6 kinases: From basic science to cancer therapy. Science (1979). 375, (2022).

3. Baker, S. J., Poulikakos, P. I., Irie, H. Y., Parekh, S. & Reddy, E. P. CDK4: a master regulator of the cell cycle and its role in cancer. Genes Cancer 13, 21–45 (2022).

4. Asciolla, J. J., Wu, X., Adamopoulos, C., Gavathiotis, E. & Poulikakos, P. I. Resistance mechanisms and therapeutic strategies of CDK4 and CDK6 kinase targeting in cancer. *Nat*. Cancer 6, 24–40 (2025).

5. Knudsen, E. S. et al. Pan-cancer molecular analysis of the RB tumor suppressor pathway. *Commun*. Biol. 3, 158 (2020).

6. Serra, S. & Chetty, R. p16. J. Clin. Pathol. 71, 853–858 (2018).

7. Thullberg, M. et al. Distinct versus redundant properties among members of the INK4 family of cyclin-dependent kinase inhibitors. FEBS Lett. 470, 161–166 (2000).

8. Shanabag, A., Armand, J., Son, E. & Yang, H. W. Targeting CDK4/6 in breast cancer. Exp. Mol. Med. 57, 312–322 (2025).

9. Wu, X. et al. Distinct CDK6 complexes determine tumor cell response to CDK4/6 inhibitors and degraders. *Nat*. Cancer 2, 429–443 (2021).

10. Li, Q. et al. INK4 Tumor Suppressor Proteins Mediate Resistance to CDK4/6 Kinase Inhibitors. Cancer Discov. 12, 356–371 (2022).

11. Morrison, L., Loibl, S. & Turner, N. C. The CDK4/6 inhibitor revolution — a game-changing era for breast cancer treatment. Nat. Rev. Clin. Oncol. 21, 89–105 (2024).

12. Palmer, C. L. et al. CDK4 selective inhibition improves preclinical anti-tumor efficacy and safety. Cancer Cell 43, 464–481.e14 (2025).

13. Gallego, G. M. et al. Discovery of Atirmociclib (PF-07220060): A Potent and Selective CDK4 Inhibitor. J. Med. Chem. 68, 26085–26098 (2025).

14. Knudsen, E. S., Witkiewicz, A. K. & Kabraji, S. The evolving landscape of CDK inhibitor use in breast cancer therapy and beyond. Nat. Rev. Drug Discov. 10.1038/s41573-026-01431-5 (2026) doi:10.1038/s41573-026-01431-5.

15. Jiang, B., et al. Development of Dual and Selective Degraders of Cyclin-Dependent Kinases 4 and 6. Angew. Chem. Int. Ed. 58, 6321–6326 (2019).

16. Marei, H. E. et al. Targeting CDKs in cancer therapy: advances in PROTACs and molecular glues. NPJ Precis. Oncol. 9, 204 (2025).

17. Oleinikovas, V., Gainza, P., Ryckmans, T., Fasching, B. & Thomä, N. H. From Thalidomide to Rational Molecular Glue Design for Targeted Protein Degradation. Annu. Rev. Pharmacol. Toxicol. 64, 291–312 (2024).

18. Békés, M., Langley, D. R. & Crews, C. M. PROTAC targeted protein degraders: the past is prologue. Nat. Rev. Drug Discov. 21, 181–200 (2022).

19. Chamberlain, P. P. & Hamann, L. G. Development of targeted protein degradation therapeutics. Nat. Chem. Biol. 15, 937–944 (2019).

20. Krönke, J. et al. Lenalidomide Causes Selective Degradation of IKZF1 and IKZF3 in Multiple Myeloma Cells. Science (1979). 343, 301–305 (2014).

21. Lu, G. et al. The Myeloma Drug Lenalidomide Promotes the Cereblon-Dependent Destruction of Ikaros Proteins. Science (1979). 343, 305–309 (2014).

22. Krönke, J. et al. Lenalidomide induces ubiquitination and degradation of CK1α in del(5q) MDS. Nature 523, 183–188 (2015).

23. Matyskiela, M. E. et al. A novel cereblon modulator recruits GSPT1 to the CRL4CRBN ubiquitin ligase. Nature 535, 252–257 (2016).

24. Sievers, Q. L. et al. Defining the human C2H2 zinc finger degrome targeted by thalidomide analogs through CRBN. Science (1979). 362, (2018).

25. Donovan, K. A. et al. Thalidomide promotes degradation of SALL4, a transcription factor implicated in Duane Radial Ray syndrome. Elife 7, (2018).

26. Steger, M. et al. Unbiased mapping of cereblon neosubstrate landscape by high-throughput proteomics. Nat. Commun. 16, 7773 (2025).

27. Annunziato, S. et al. Cereblon induces G3BP2 neosubstrate degradation using molecular surface mimicry. Nat. Struct. Mol. Biol. 33, 479–487 (2026).

28. Baek, K. et al. Unveiling the hidden interactome of CRBN molecular glues. Nat. Commun. 16, 6831 (2025).

29. Caldwell, J. J. et al. Developing potent and selective TBK1 molecular glue degraders for cancer immunotherapy. Preprint at 10.64898/2026.01.30.702304 (2026).

30. Shashikadze, B. et al. Integrated proteomic screening reveals design principles of CRBN molecular glue degraders. bioRxiv Preprint at 10.64898/2026.03.08.710269 (2026).

31. Razumkov, H. et al. Discovery of CRBN-Dependent WEE1 Molecular Glue Degraders from a Multicomponent Combinatorial Library. J. Am. Chem. Soc. 146, 31433–31443 (2024).

32. Petzold, G. et al. Mining the CRBN target space redefines rules for molecular glue–induced neosubstrate recognition. Science (1979). 389, (2025).

33. Gkountela, S., et al. Abstract LB422: Selective targeting of CDK2 using molecular glue degraders for the treatment of HR-positive/HER2-negative breast cancer. Cancer Res. 85, LB422–LB422 (2025).

34. Min, J., et al. Phenyl-Glutarimides: Alternative Cereblon Binders for the Design of PROTACs. Angew. Chem. Int. Ed. 60, 26663–26670 (2021).

35. Sperling, A. S. et al. Patterns of substrate affinity, competition, and degradation kinetics underlie biological activity of thalidomide analogs. Blood 134, 160–170 (2019).

36. Lu, D. et al. CREPT Accelerates Tumorigenesis by Regulating the Transcription of Cell-Cycle-Related Genes. Cancer Cell 21, 92–104 (2012).

37. Li, M., Ma, D. & Chang, Z. Current understanding of CREPT and p15RS, carboxy-terminal domain (CTD)-interacting proteins, in human cancers. Oncogene 40, 705–716 (2021).

38. Steger, M. et al. Time-resolved in vivo ubiquitinome profiling by DIA-MS reveals USP7 targets on a proteome-wide scale. Nat. Commun. 12, 5399 (2021).

39. Amako, Y. et al. The contribution of cyclic imide stereoisomers on cereblon-dependent activity. Chem. Sci. 16, 11519–11529 (2025).

40. Reist, M., Carrupt, P.-A., Francotte, E. & Testa, B. Chiral Inversion and Hydrolysis of Thalidomide: Mechanisms and Catalysis by Bases and Serum Albumin, and Chiral Stability of Teratogenic Metabolites. Chem. Res. Toxicol. 11, 1521–1528 (1998).

41. Zhou, Y. et al. Heteroaryl Glutarimides and Dihydrouracils as Cereblon Ligand Scaffolds for Molecular Glue Degrader Discovery. ACS Med. Chem. Lett. 15, 2158–2163 (2024).

42. Scattolin, D. et al. Clinical insight on the pathway of SLFN11 as emergent biomarker in SCLC. Crit. Rev. Oncol. Hematol. 215, 104901 (2025).

43. Kroupova, A. et al. Design of a Cereblon construct for crystallographic and biophysical studies of protein degraders. Nat. Commun. 15, 8885 (2024).

44. Sang, N., Caro, J. & Giordano, A. Adenoviral E1A: everlasting tool, versatile applications, continuous contributions and new hypotheses. Front. Biosci. 7, d407–13 (2002).

45. Kamb, A. et al. A Cell Cycle Regulator Potentially Involved in Genesis of Many Tumor Types. Science (1979). 264, 436–440 (1994).

46. Arafeh, R., Shibue, T., Dempster, J. M., Hahn, W. C. & Vazquez, F. The present and future of the Cancer Dependency Map. Nat. Rev. Cancer 25, 59–73 (2025).

47. Liberzon, A. et al. The Molecular Signatures Database Hallmark Gene Set Collection. Cell Syst. 1, 417–425 (2015).

48. Pack, L. R., Daigh, L. H., Chung, M. & Meyer, T. Clinical CDK4/6 inhibitors induce selective and immediate dissociation of p21 from cyclin D-CDK4 to inhibit CDK2. Nat. Commun. 12, 3356 (2021).

49. Russo, A. A., Tong, L., Lee, J.-O., Jeffrey, P. D. & Pavletich, N. P. Structural basis for inhibition of the cyclin-dependent kinase Cdk6 by the tumour suppressor p16INK4a. Nature 395, 237–243 (1998).

50. Glennie, L. et al. The Contribution of Native Protein Complexes to Targeted Protein Degradation. ACS Chem. Biol. 21, 1095–1111 (2026).

51. Gump, J., Stokoe, D. & McCormick, F. Phosphorylation of p16 Correlates with Cdk4 Association. Journal of Biological Chemistry 278, 6619–6622 (2003).

52. Lu, Y., Ma, W., Li, Z., Lu, J. & Wang, X. The interplay between p16 serine phosphorylation and arginine methylation determines its function in modulating cellular apoptosis and senescence. Sci. Rep. 7, 41390 (2017).

53. Ben-Saadon, R. et al. The Tumor Suppressor Protein p16 and the Human Papillomavirus Oncoprotein-58 E7 Are Naturally Occurring Lysine-less Proteins That Are Degraded by the Ubiquitin System. Journal of Biological Chemistry 279, 41414–41421 (2004).

54. Ragione, F. Della et al. Biochemical Characterization of p16 - and p18-containing Complexes in Human Cell Lines. Journal of Biological Chemistry 271, 15942–15949 (1996).

55. Lipinski, C. A., Lombardo, F., Dominy, B. W. & Feeney, P. J. Experimental and computational approaches to estimate solubility and permeability in drug discovery and development settings 1PII of original article: S0169-409X(96)00423-1. The article was originally published in Advanced Drug Delivery Reviews 23 (1997) 3–25. 1. Adv. Drug Deliv. Rev. 46, 3–26 (2001).

56. Rappsilber, J., Ishihama, Y. & Mann, M. Stop and Go Extraction Tips for Matrix-Assisted Laser Desorption/Ionization, Nanoelectrospray, and LC/MS Sample Pretreatment in Proteomics. Anal. Chem. 75, 663–670 (2003).

57. Brown, L. Y., Dong, W. & Kantor, B. An Improved Protocol for the Production of Lentiviral Vectors. STAR Protoc. 1, 100152 (2020).

58. Meier, F. et al. diaPASEF: parallel accumulation–serial fragmentation combined with data-independent acquisition. Nat. Methods 10.1038/s41592-020-00998-0 (2020) doi:10.1038/s41592-020-00998-0.

59. Szyrwiel, L., Sinn, L., Ralser, M. & Demichev, V. Slice-PASEF: fragmenting all ions for maximum sensitivity in proteomics. bioRxiv 2022.10.31.514544 (2022) doi:10.1101/2022.10.31.514544.

60. Skowronek, P., Wallmann, G., Wahle, M., Willems, S. & Mann, M. An accessible workflow for high-sensitivity proteomics using parallel accumulation–serial fragmentation (PASEF). Nat. Protoc. 20, 1700–1729 (2025).

61. Demichev, V., Messner, C. B., Vernardis, S. I., Lilley, K. S. & Ralser, M. DIA-NN: neural networks and interference correction enable deep proteome coverage in high throughput. Nat. Methods 17, 41–44 (2020).

62. Ritchie, M. E. et al. limma powers differential expression analyses for RNA-sequencing and microarray studies. Nucleic Acids Res. 43, e47–e47 (2015).

